# RPRD Proteins Control Transcription in Human Cells

**DOI:** 10.1101/2021.06.20.449126

**Authors:** Kinga Winczura, Hurmuz Ceylan, Monika Sledziowska, Matt Jones, Holly Fagarasan, Jianming Wang, Marco Saponaro, Roland Arnold, Daniel Hebenstreit, Pawel Grzechnik

## Abstract

The regulation of transcription is an essential process that allows the cell to respond to various internal and external signals. RNA Polymerase II (Pol II) activity is controlled by a number of factors which bind to the C-terminal domain (CTD) of its largest subunit, RPB1, and stimulate or suppress RNA synthesis. Here, we demonstrate that CTD-interacting proteins, RPRD2, RPRD1B and RPRD1A act as negative regulators of transcription and their levels inversely correlate with the accumulation of nascent and newly transcribed RNA in human cells. We show that the RPRD proteins form mutually exclusive complexes with Pol II to coordinate their roles in transcriptional control. Our data indicate that RPRD2 exerts the most substantial impact on transcription and has the potential to alter key biological processes including the cellular stress response and cell growth.

## INTRODUCTION

The regulation of gene expression is an indispensable process required for proper functioning and survival of the cell. Both activation and suppression of gene expression play equally important roles that allow cells to adjust their response to various internal and external stimuli. The downregulation of transcription facilitates the silencing or reduction of gene expression by limiting the level of RNA synthesis. The presence of RNA Polymerase II (Pol II) on a gene that is not highly expressed, allows for a rapid shift from an “off” to “on” state when it is required (Adelman et al., 2009; Core and Adelman, 2019; Danko et al., 2013; Gilchrist et al., 2012). Such transitions are essential for survival strategies including stress and cellular signalling (de Nadal et al., 2011). Once Pol II is recruited to the gene promoter, reduced RNA synthesis can be achieved by premature transcription termination, decreased transcription rates or increased Pol II pausing on the promoter (Jonkers et al., 2014; Kamieniarz-Gdula et al., 2019).

In higher eukaryotes, Pol II initiates RNA synthesis and pauses 30-60 nucleotides from the initiation site (Adelman and Lis, 2012). This prevalent rate-limiting event called promoter-proximal pausing provides a point of regulation that integrates multiple signals that decide if transcription progresses to the elongation phase. This step is associated with specific phosphorylation of the Pol II C- terminal domain (CTD) and the exchange of factors on the Pol II complex. The human CTD consists of 52 heptapeptide repeats (consensus Y1-S2-P3-T4-S5-P6-S7) and phosphorylation of Y1, S2, T4, S5 and S7 is associated with specific stages of the transcription cycle (Buratowski, 2003; Eick and Geyer, 2013; Hsin and Manley, 2012). Initiated Pol II, phosphorylated at S5 of the CTD, starts transcribing towards the +1 nucleosome where it becomes stabilised by negative elongation factor (NELF) and DRB-sensitivity-inducing factor (DSIF) (Adelman and Lis, 2012; Kwak et al., 2013). The positive transcription elongation factor-b (P-TEFb) coordinates release from pausing and the transition into the processive phase. P-TEFb phosphorylates NELF, DSIF and Pol II CTD at S2 (Peterlin and Price, 2006). Simultaneously, S5 residues become dephosphorylated by RPAP2 (Egloff et al., 2012; Mosley et al., 2009). NELF dissociates from the Pol II complex, while DSIF converts into a positive elongation factor (Peterlin and Price, 2006). The early stages of transcription and the elongation phase are supported by a conserved Paf1 complex (Paf1C), which can both stimulate and inhibit Pol II transition from promoter-proximal pausing and is also required for establishing boundaries between heterochromatin (Francette et al., 2021; Hou et al., 2019; Krogan et al., 2003; Verrier et al., 2015). An alternative mechanism of transition from initiation to elongation is mediated by a substoichiometric Pol II subunit GDOWN1 that excludes TFIIF and TFIIB initiation factors from promoter-bound Pol II leading to stable promoter-proximal pausing (Cheng et al., 2012; Jishage et al., 2012; Jishage et al., 2018). The transition to elongation requires mediator to overcome the inhibitory effect of GDOWN1 (Cheng et al., 2012).

All steps in the transcriptional cycle are supported by factors interacting with the CTD of Pol II. These factors act to coordinate co-transcriptional enzymatic activities in response to Pol II CTD phosphorylation status and/or signals in the nascent RNA (Hsin and Manley, 2012). One superfamily of these interactors contains proteins with the CTD-interacting domains (CID) that recognise specific CTD phosphorylation patterns. These essential factors include PCF11, SCAF4 and SCAF8 which are involved in the regulation of transcription termination and the formation of mRNAs 3’ ends (Gregersen et al., 2019; Kamieniarz-Gdula et al., 2019). The three RPRD (Regulation of nuclear pre-mRNA domain) proteins, RPRD1A, RPRD1B and RPRD2 (RPRDs) also belong to this superfamily (Ni et al., 2011). RPRD1A and RPRD1B are paralogs (80% similarity) and, along with the CID, contain coiled-coil (CC) domains (Figure S1A) (Ni et al., 2011; Ni et al., 2014), that mediate their interaction. Both proteins bind to the CTD phosphorylated at S5 and have been shown to act together in the recruitment of RPAP2 and HDAC1 which act as CTD S5 phosphatase and CTD deacetylase, respectively in the early stages of transcription (Ali et al., 2019; Ni et al., 2014). Therefore, the RPRD1A-RPRD1B heterodimer has been suggested to promote transcription (Ali et al., 2019; Ni et al., 2014). Despite this fact, there is a discrepancy in the phenotypes associated with cellular levels of either RPRD1B or RPRD1A. Increased expression of RPRD1B (alias CREPT: cell cycle-related and expression elevated protein in tumour) stimulates cell proliferation and is associated with carcinogenesis (Lu et al., 2012). In contrast, RPRD1A has been identified as a tumour suppressor and its increased expression slows cellular growth (Liu et al., 2002). The third protein from the RPRD family, RPRD2, remains functionally uncharacterised. RPRD2 is much larger than its two counterparts but its N-terminus shares homology with RPRD1A and RPRD1B (Ni et al., 2014). Apart from the CID, it contains two other domains enriched in serines and prolines (Ser- and Pro-rich, respectively) (Ni et al., 2011). Recently, RPRD2 has been identified as a component in a pathway that suppresses the replication of human immunodeficiency virus HIV-1 (Gibbons et al., 2020; Marno et al., 2014).

Apart from biochemical and functional association of RPRD1B and RPAP2 in promoting transcription, RPRD1B has been reported to co-purify with negative transcription regulator GDOWN1 (Cheng et al., 2012; Jishage et al., 2012; Ni et al., 2011). Moreover, both RPRD1B and RPRD2 co-purify with DSIF and PAF1C (Hein et al., 2015; Hou et al., 2019). These interactions suggest that RPRD proteins may have additional roles in transcriptional regulation. Here, we demonstrate that RPRDs exert a negative impact on transcription in human cells. By employing RNA metabolic labelling with thiouridine (4sU) followed by sequencing (4sU-seq), we show that the depletion of the RPRD proteins resulted in increased nascent RNA production, whereas their overexpression lead to downregulation of transcription. These transcriptional changes varied depending on which RPRD protein was deregulated. We show that the most effective transcriptional suppressor among the RPRD proteins is RPRD2. Our data indicate its potential to regulate the cell cycle progression, cell growth as well as the cellular stress response, in particular to viral infections. The use of CRISPR-engineered cell lines expressing FLAG tagged RPRDs allowed us to apply a standardized affinity capture protocol for all three RPRDs. This enabled direct comparison of the individual RPRDs interactomes. We show that RPRD1B forms mutually exclusive complexes with RPRD1A and RPRD2, potentially coordinating their roles in the transcription control. Altogether, we reveal a novel role for the RPRD proteins in the downregulation of gene expression.

## RESULTS

### Depletions of the RPRD proteins increase RNA synthesis

Differential RPRD1B and RPRD1A protein levels have been associated with many human tumours (Li et al., 2021; Lu et al., 2012; Wu et al., 2010). This indicates that increased or decreased expression of RPRDs plays an important role in the regulation of transcription and gene expression. Thus, in our experimental approach, we aimed to explore the effects of RPRD1B, RPRD1A and RPRD2 depletion and overexpression on the nascent transcriptome (Figure S1B). Depletions of the RPRDs (KD) were achieved by treating HEK293T cells with siRNA, while to overexpress (OE) the RPRDs, we used the Flp-In T-REx 293 system, described below.

First, we investigated how the decreased levels of individual RPRDs affect RNA synthesis. We depleted RPRD2, RPRD1B or RPRD1A for 48 hours (Figure S1C) and employed RNA metabolic labelling with 4sU and sequenced the isolated labelled RNA (Figure 1A). In this approach, we incubated cells with 4sU for 5 minutes, thus the levels of labelled RNA reflected nascent and newly synthesised RNAs (nnsRNAs) which was indicated by a decreased ratio of reads from exons and introns. Total RNA isolated from HEK293T cells was mixed (10:1) with RNA from yeast *Saccharomyces cerevisiae* labelled for 15 minutes with thiouracil (4tU). This spike-in was later used for normalisation of the nnsRNA levels in each sample to compensate for differences introduced by the variable RNA purification. The efficiency of nnsRNA precipitation was confirmed by qPCR (Figure S1D). Sequenced RNA reads were aligned to the human genome, normalised to the amount of co-purified spike-in yeast RNA and counted to obtain the expression levels of each of ~60,000 transcription units (TUs). The normalization was performed according to DEseq’2 median ratio method, which accounts for the sequencing depth and RNA composition (Anders and Huber, 2010). Next, we compared each KD sample against the control. The accumulation of nnsRNA for individual genes in KD and control samples were presented on scatter plots (Figure 1B). The accumulation of the nnsRNAs depicted above the zero-difference line, indicates they were transcribed to a higher extent. The depletions of individual RPRD proteins resulted in increased levels of nnsRNA when compared to the control. RPRD2 KD displayed the most pronounced effect and RPRD1A KD affected the transcriptome the least, as demonstrated by the relation of the regression line to the zero-line (Figure 1B). We observed a positive fold change (FC) for most of the genes in KD samples, when compared to the control, confirming globally enhanced transcription caused by the RPRDs depletions (Figure 1C).

**Figure 1.**
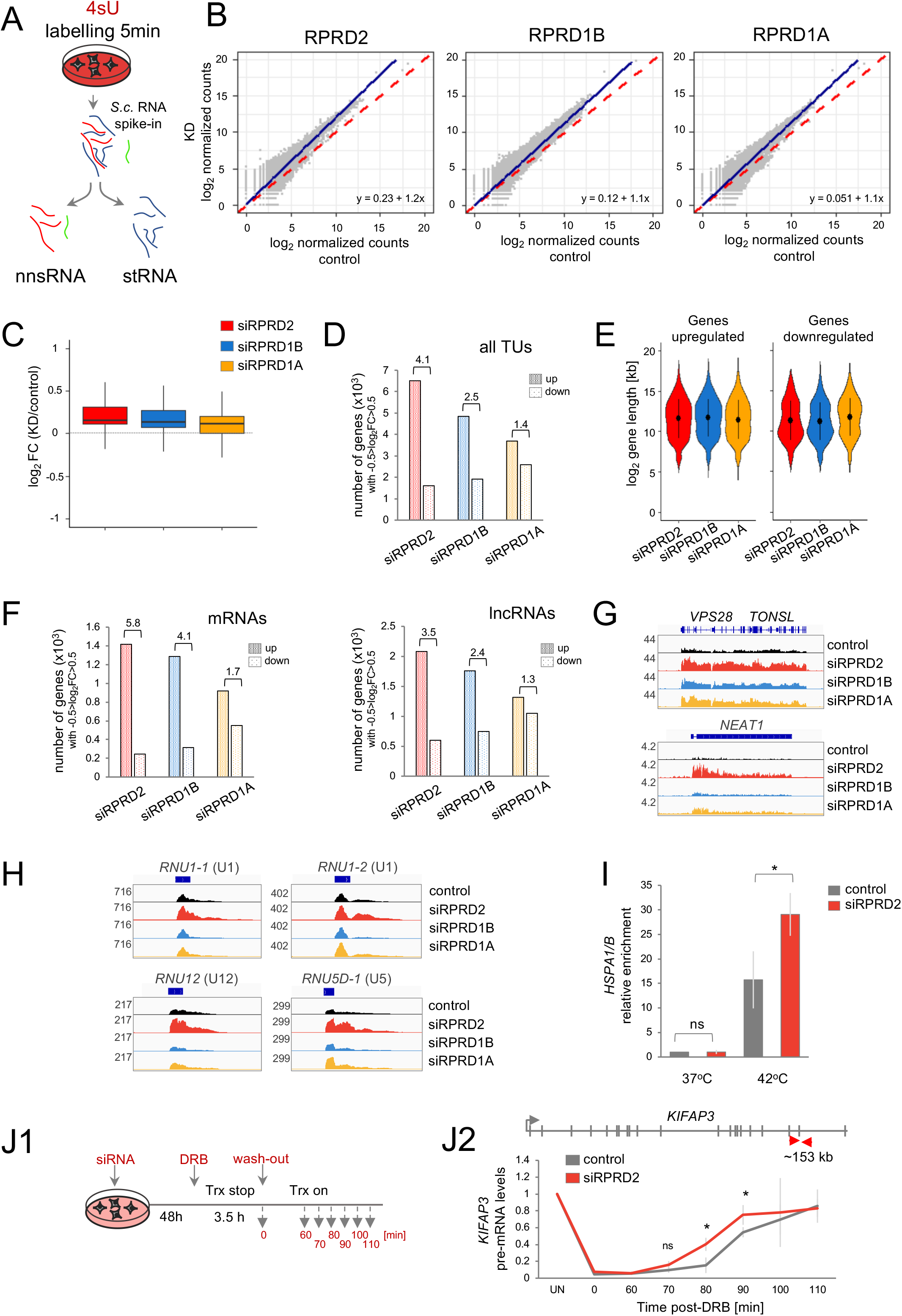
The impact of the RPRDs depletion on RNA synthesis. **A.** Isolation of the nascent and newly synthesised RNA (nnsRNA) from the steady-state RNA (stRNA) fraction. Cells were harvested after a 5 min incubation with 4sU. Isolated total RNA was spiked-in with 4tU-labelled RNA from *S. cerevisiae* and biotinylated. nnsRNA was separated from the stRNA fraction on streptavidin beads. **B.** Levels of nnsRNAs upon RPRDs knockdown (KD), 4sU-seq analysis. Scatter plot representing the values of *log_2_* normalised counts for each gene in the KD samples (y-axis) against the control samples (x-axis). The blue lines represent the regression trends, the equations of which are shown in the corner of each plot. The dashed red diagonals denote 0 *log_2_* fold change (FC). **C**. Box plots showing the *log_2_* FC in nnsRNA accumulation for RPRDs KD samples over control. **D**. Bar plots presenting the number of genes displaying the *log_2_* FC greater than 0.5 (up, densely dotted) or lower than −0.5 (down, lightly dotted) in 4sU-seq analysis for each KD sample. The numbers above the bars indicate the ratio between upregulated and downregulated genes for each KD sample. **E**. Violin plots showing the length-dependent distribution of upregulated and downregulated genes in each KD sample. **F.** Number of mRNAs and lncRNAs showing *log_2_* FC greater than 0.5 or lower than −0.5 in 4sU-seq analysis. The numbers above the bars indicates the ratio between upregulated and downregulated genes **G**. IGV (Integrative Genomics Viewer) tracks showing the accumulation of nnsRNA levels for *VPS28* and *TONSL* mRNA (top) and *NEAT1* lncRNA (bottom) in KD samples and the control. The tracks show counts x10^5^. **H.** IGV tracks showing the accumulation of snRNAs in RPRDs KD samples (counts x10^5^). **I.** RPRD2 KD increased expression of heat-shocked induced *HSP1A/B* gene; RT-qPCR analysis. A bar plot showing the relative enrichment of *HSP1A/B* mRNA in control (grey) and RPRD2 KD (red) in unconditioned cells (37°C) and upon the heat shock (42°C). Error bars represent the standard deviation calculated from three experiments. An asterisk (*) denotes p-value < 0.05; ns – non-significant. **J1.** A diagram showing an experimental approach to study Pol II transcriptional rates. Cells were treated with DRB for 3.5h to stop transcription (Trx). Samples were collected after DRB removal at the time points indicated on the diagram. **J2.** RPRD2 KD decreased the transcription rate over *KIFAP3* gene; RT-qPCR analysis. *Top* A schematic representation showing introns and exons of the *KIFAP3* gene. Red arrows point to the position of the amplicon tested in the analysis. *Bottom* A graph showing the relative levels of *KIFAP3* pre-mRNA (y-axis) at each time point (x-axis) normalized to untreated cells (UN) for control (grey) and RPRD2 KD (red). Error bars represent the standard deviation calculated from three experiments. An asterisk (*) denotes p-value < 0.05; ns – non-significant.

Next, we selected the TUs whose levels were most affected by the KDs (−0.5 > log2 FC (KD/control) > 0.5) and discarded outliers (Figure 1D). We found that RPRD2 depletion increased transcription of 6511 and decreased 1607 TUs (ratio up/down = 4.1), while RPRD1B depletion increased levels of 4846 and decreased levels of 1916 TUs (ratio = 2.5). The number of positively and negatively affected TUs in RPRD1A KD was 3693 and 2589 respectively, resulting in the smallest ratio (1.4). The size of upregulated TUs did not differ between the samples with the median length ~3.5 kb (Figure 1E). We also tested how RPRDs affected RNA synthesis of different RNA classes, mRNAs, long non-coding RNAs (lncRNAs) and small nuclear RNAs (snRNAs). In the case of mRNA (Figure 1F, left panel), the ratios of increased to decreased mRNAs was 5.8 for RPRD2 KD and 4.1 for RPRD1B KD, in both cases confirming general gene upregulation. RPRD1A KD resulted in the lower ratio of increased to decreased mRNAs (1.7). A similar effect of RPRD KDs was observed for lncRNAs (Figure 1F, right panel). Analysis of single gene examples visualised as reads tracks in Integrated Genome Viewer (IGV) confirmed these observations. Depletion of RPRDs resulted in altered levels of nnsRNA for Pol II-transcribed protein-coding genes *VPS28, TONSL* and *KPNB1* as well as lncRNA *NEAT1* but not Pol I-transcribed rRNA (Figure 1G and S1E). Due to low number of independently transcribed snRNA genes we inspected their accumulation by analysing the 4sU-seq reads tracks in the genome browser (Figure 1H). We found that in most cases only RPRD2 depletion increased accumulation of snRNAs as it is seen for U1, U5 and U12. Altogether, this data indicates that RPRD2 and RPRD1B depletion leads to the upregulation of RNA synthesis, while RPRD1A KD has a nebulous effect on transcription.

To test the accumulation of steady-state RNA in RPRDs KD, we performed 3’RNA sequencing (3’RNA-seq) on total RNA. Effects of depletions on steady-state RNAs were much less evident than for nnsRNAs, with only a small fraction of transcripts being deregulated (Figure S1F), which is in line with previous reports (Ali et al., 2019). This may be a result of the cells ability to adjust RNA levels by inhibiting or enhancing RNA degradation pathways (Haimovich et al., 2013; Sun et al., 2013). Alternatively, the most erroneous changes introduced by the depletions of RPRDs need more time to accumulate than the 48h we applied in our experiments. A more direct effect of transcription on steady-state RNAs can be envisaged by analysis of the accumulation of inducible genes.

Since RPRD2 KD exhibited the strongest effect on transcription, by globally upregulating nnsRNA levels, we tested if its depletion affects the expression of inducible genes, such as stress responsive *HSPA1A/B*. We applied heat-shock to stress-sensitive MEF cells (Vihervaara et al., 2021), by transferring them from 37°C to 42°C for 1 hour and assessing the changes in *HSPA1A/B* mRNA accumulation by qPCR analysis. Our data showed that upon stress induction, the *HSPA1A/B* level almost doubled in RPRD2 KD compared to control but remained unchanged in unconditioned cells (Figure 1I). This finding further substantiates a role for RPRD2 in transcription downregulation.

Next, we investigated if the increase in nnsRNA levels upon RPRDs KD could be attributed to changes in the speed of transcription. To test Pol II transcription rates upon RPRD2 depletion, we performed a “DRB stop-chase” experiment (Figure J1) (Saponaro et al., 2014). We treated cells with 5,6-dichloro-1-β-d-ribofuranosylbenzimidazole (DRB) for 3.5 hours to inhibits CDK9 activity, thereby arresting Pol II in promotor-proximal paused state and blocking elongation. After this time, DRB was removed from the medium to allow the restoration of transcription. Cells were collected at experimentally established time points (0 and 60-110 min) and transcription rates were assessed by qPCR analysis. To enrich for nnsRNA synthesised during the release phase, we used an amplicon spanning over the exon-intron 20 junction of the *KIFAP3* gene, which is located ~150 kb from the promoter. The *KIFAP3* pre-mRNA started to accumulate in the control sample at ~80 minutes after DRB retraction, indicating that Pol II required 80 minutes to transcribe 150 kb of this gene. For the RPRD2 KD, the presence of the signal from exon-intron 20 was accelerated by ~10 minutes. Therefore, this assay revealed that upon depletion of RPRD2, *KIFAP3* is transcribed faster than in the control. We concluded that affected Pol II transcription rates resulted in an increased accumulation of 4sU labelled nnsRNA observed upon RPRDs KD.

### Overexpression of the RPRD proteins decreases RNA synthesis

Since RPRD1B increased levels are associated with faster proliferation (Lu et al., 2012), and RPRD1B and RPRD2 are often found to be upregulated in cancer (Figure S2A) (Tate et al., 2019), we also explored the effects of the elevated levels of RPRD2, RPRD1B and RPRD1A on transcription. To overexpress RPRDs we used the Flp-In T-REx 293 system. We stably integrated the coding sequences of C-terminally 3xFLAG (3F)-tagged RPRD1B, RPRD1A or RPRD2 into an FRT locus of this commercially engineered HEK293 cell line and expressed them from the tetracycline (Tet)-inducible promoter (Figure S2B). Each tagged cell line also expressed endogenous RPRDs from the unaffected genomic loci. We empirically established the Tet concentration (50 ng/ml) necessary to achieve high OE (Figure S2C). To test the effects of individual RPRDs OE on the transcriptome, we employed a similar approach as was used for the KD experiments. We performed 4sU-seq analysis after inducing OE of RPRDs and control GFP for 48 hours. Isolated RNA was spiked-in with *S. cerevisiae* RNA and the purified labelled fraction was sequenced. We employed the same bioinformatics pipeline as for RPRD KD experiments. First, we analysed each OE sample against the control overexpressing GFP and represented the results on scatter plots (Figure 2A). The OE of RPRD2 and RPRD1B resulted in decreased levels of nnsRNA when compared to the control, while RPRD1A OE moderately affected the transcriptome. The decreased average FC of nnsRNAs in the OEs samples when compared to control (Figure 2B) confirmed the downregulatory effects of RPRD2 and RPRD1B OE on transcription. Interestingly, in contrast to RPRD2 and RPRD1B, the upregulation of nnsRNA levels by RPRD1A OE was more dominant than transcription repression. Next, we selected the TUs whose levels were most affected by the OE (−0.5 > log2 FC (OE/control) > 0.5) (Figure 2C). This analysis revealed that RPRD2 OE increased transcription of 1492 and decreased transcription of 5258 TUs (ratio up/down = 0.35), while RPRD1B OE increased 1916 and decreased levels of 3345 TUs (ratio = 0.6). In the case of RPRD1A, the number of TUs that were upregulated or downregulated, were 3860 and 2033, respectively, resulting in the ratio 1.9. The size of downregulated genes for RPRD2 and RPRD1B OE was similar (median length ~10kb) but distinct to RPRD1A OE (median 4.5kb) (Figure 2D). We also tested how RPRDs OE affected RNA synthesis of various RNA classes from this data set (Figure 2E). RPRD2 and RPRD1B OE mostly decreased nnsRNAs of protein-coding genes. This is shown as ratio (increased/decreased) of 0.2 for RPRD2 OE and 0.5 for RPRD1B OE. RPRD1A OE increased mRNAs synthesis resulting in the ratio of 2.3. RPRDs OE affected the transcription of lncRNAs in a similar way to protein-coding genes (Figure 2E, right panel). Single gene examples visualised in the IGV genome viewer confirmed these results. Depletion of RPRD2 and RPRD1B resulted in a severe decrease of nnsRNA for protein-coding *MYC, VPS28 and TONSL*, and lncRNA *XIST* but not Pol I-transcribed rRNA (Figure 2F and S2D). snRNA transcription was the least affected by the RPRDs OE (Figure 2G). In the case of U5 we observed an opposite effect of RPRD2 OE to RPRD1B and RPRD1A OE.

**Figure 2.**
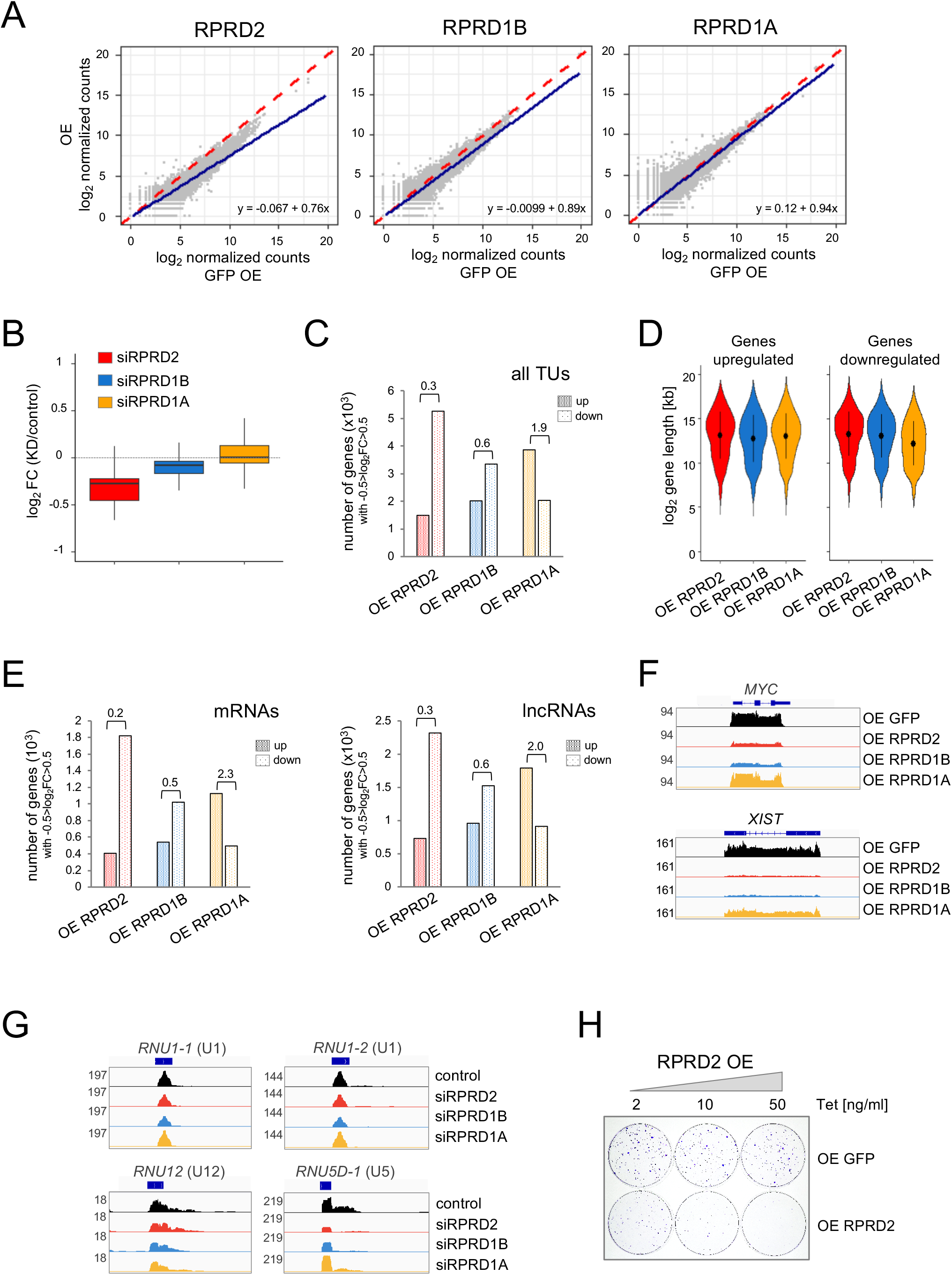
The impact of the RPRDs overexpression on RNA synthesis. **A.** Levels of nnsRNAs upon RPRDs overexpression (OE), 4sU-seq analysis. Scatter plots representing the values of *log_2_* normalised counts for each gene in the OE samples (y-axis) against the control samples (x-axis). The blue lines represent the regression trends, the equations of which are shown in the corner of each plot. The dashed red diagonals denote 0 *log_2_* fold change (FC). **B**. Box plots showing the *log_2_* FC in nnsRNA accumulation for RPRDs OE samples over control. **C**. Bar plots presenting the number of genes displaying the *log_2_* FC greater than 0.5 (up, densely dotted) or lower than −0.5 (down, lightly dotted) in 4sU-seq analysis for each OE sample. The numbers above the bars indicate the ratio between upregulated and downregulated genes for each OE sample. **D**. Violin plots showing the length-dependent distribution of upregulated and downregulated genes in each OE sample. **E.** Number of mRNAs and lncRNAs showing the *log_2_* FC greater than 0.5 or lower than −0.5 in 4sU-seq analysis. The numbers above the bars indicates the ratio between upregulated and downregulated genes **F**. IGV (Integrative Genomics Viewer) tracks showing the accumulation of nnsRNA levels for *MYC* mRNA (top) and *XIST* lncRNA (bottom) in OE samples and the control.The tracks show counts x10^5^. **G.** IGV tracks showing the accumulation of snRNAs in RPRDs OE samples (counts x10^5^). **H.** Colony forming unit assay showing the growth of the cells overexpressing RPRD2. Cells growing on Tet-free medium were seeded 100 cells per well and RPRD2 OE was induced with a different concentration of Tet. After 9 days cells were fixed and stained with crystal violet.

This suggests that short genes were less susceptible to the inhibitory effects of the RPRDs on transcription. Next, we tested if transcription inhibition observed upon RPRD2 OE had an effect on cell proliferation (Figure 2H). We induced OE of either GFP or RPRD2 with variable concentration of Tet for 9 days. Even a relatively small increase in RPRD2 constitutive expression (induced with 2 ng/ml Tet) affected the cell growth when compared to cells OE GFP. Higher OE of RPRD2 almost completely abolished cell growth. Altogether, our data demonstrated that the RPRD proteins, in particular RPRD2, act as general negative regulators of transcription.

### The RPRDs interactomes

To understand how RPRDs control transcription, we decided to look at their interactions with other proteins. RPRDs were reported to co-purify with a range of transcriptional factors, including RECQL5, GDOWN1, RPAP2 and PAF1 complex (Hein et al., 2015; Ni et al., 2011); however, their interactomes had not been directly compared. To that end, we performed immunoprecipitation (IP) of all RPRDs followed by mass spectrometry (MS) analysis. To avoid differences in IP profiles caused by different efficiency and specificity of commercially available antibodies, we inserted a 3F tag at the C-terminus of each RPRD in HEK293T cells using the CRIPSR-Cas9 approach. To ensure that both alleles were tagged, we transfected cells with two donor plasmids harbouring the 3F sequence in frame with the last exon of the gene, followed by either blasticidin or hygromycin resistance genes (Figure 3A). Single clones growing on selective medium containing blasticidin and hygromycin were tested by western blot for the presence of the tagged protein (Figure S3A). Next, we used anti-FLAG antibody to IP the tagged proteins and their interactors from cell extracts prepared in the high salt buffer (600 mM NaCl). Such stringent conditions help to enrich for strong interactions and reduce weak or unspecific ones. Immunoprecipitated fractions were analysed by MS. The results were filtered and common contaminants (Mellacheruvu et al., 2013) were discarded to reveal proteins interacting with each RPRD (Figure 2B and Table S1). All three RPRDs co-purified multiple subunits of the Pol II complex, including GDOWN1. This interaction was previously described for RPRD1B and RPRD1A (Ni et al., 2014) but not for RPRD2. Moreover, RPRD1B and RPRD1A both co-immunoprecipitated RPAP2, that was not present in the fraction associated with RPRD2. Most interestingly, we found that RPRD1B co-purified RPRD1A and RPRD2; however, both RPRD1A and RPRD2 co-purified only RPRD1B but not each other (Figure 3B). This indicates that RPRD1B forms mutually exclusive interactions either with RPRD1A or with RPRD2. Among other proteins associated with RPRD1B and RPRD1A were the BRCA2 interactor HMG20B that is involved in chromatin reorganization through histone deacetylation (Hakimi et al., 2002; Marmorstein et al., 2001); GPN-loop GTPase1, required for Pol II import to the nucleus and its assembly with regulating factors (Carre and Shiekhattar, 2011); and IMPDH, a regulator of the intracellular guanine nucleotide pool, important for signal transduction and processes involved in cellular proliferation (Gu et al., 2000). RPRD2 exclusive interactors included proteins involved in immune processes such as interferon-induced very large GTPase 1 GVINP1 and E3 ubiquitin-protein ligase TRIM21. RPRD2 also co-purified two chaperones, HSPA1 and HSPA8. Different binding partners identified in this analysis indicate an additional discrepancy between RPRD1A-RPRD1B and RPRD2 possible functions, as RPRD1A and RPRD1B co-purified mainly transcription factors, while the proteins associated with RPRD2 are potentially involved in antiviral and stress response. Further analysis of the RPRDs interactomes on SDS-PAGE revealed only few visible bands indicating that most of the interacting partners purified in substoichiometric amounts (Figure 3C). The MS analysis of these individual gel bands showed that RPRDs interact mainly with other RPRDs and with the two largest Pol II subunits, CTD-containing RPB1 and RPB2 (Table S2). Moreover, RPRD2 major interactors were HSPA1 and HSPA8 proteins. As we applied a stringent IP buffer in our experimental setup, we wondered whether more interactors could be preserved in milder conditions. For that, we applied the same IP protocol but using buffers containing increasing salt concentrations ranging from 100 mM to 1 M NaCl and analysed the interaction profiles on SDS-PAGE (Figure 3D). RPRDs co-purified more proteins in low-salt conditions which gradually disappeared with increasing salt concentration to 400 mM NaCl. Strikingly, RPRD1B-RPRD1A-Pol II and RPRD1B-RPRD2-HSP-Pol II interactions were maintained even in 0.8 and 1M NaCl. Addition of 2M urea in the IP buffer caused detachment only of the RPB2 subunit in the case of RPRD1B and RPRD1A, and RPB2 and the two chaperones in the case of RPRD2, leaving the main complex of RPRDs with RPB1 intact (Figure S3B).

**Figure 3.**
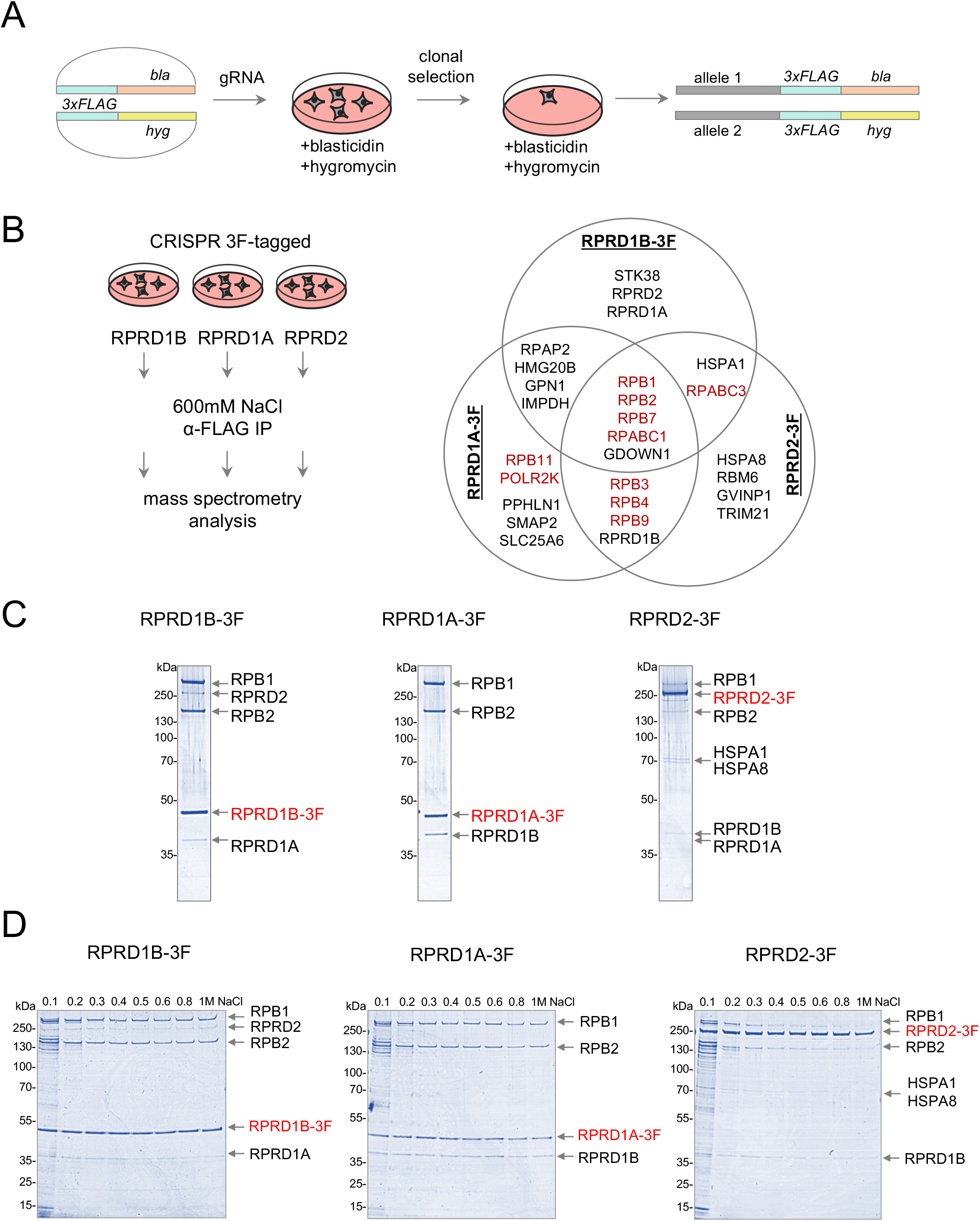
The interactome of RPRD proteins. **A.** The strategy used for CRISPR-Cas9 mediated C-terminal 3xFLAG (3F)-tagging of endogenous RPRD alleles in HEK293T cells. Two plasmids harbouring 3F tag followed by a selection marker (providing the resistance to blasticidin (bla) or hygromycin (hyg)) were co-transfected together with a gRNA vector. After transfection, cells were subjected to clonal selection by addition of blasticidin and hygromycin to the media. Single clones were propagated and tested for the presence of the tagged loci. **B.** *Left* Experimental approach to identify protein interactors of RPRDs. CRISPR cell lines expressing 3F-tagged RPRDs were used for preparation of cell extracts in a high salt buffer (600 mM NaCl) and immunoprecipitation (IP) of proteins with anti-FLAG antibodies. The IP fractions were then subjected to mass spectrometry (MS) analysis. *Right* Venn diagram showing selected common and exclusive interactors for RPRD1B, RPRD1A and RPRD2 identified in MS analysis. The subunits of Pol II are denoted in red. **C**. A Coomassie stained SDS-PAGE gel analysis of proteins co-purified with 3F-tagged RPRDs. The bait proteins (red), as well as major interactors, are indicated with arrows. The identity of the indicated bands was established by MS analysis. Protein marker sizes are shown on the left. **D.** A Coomassie stained SDS-PAGE gel analysis of proteins co-purified with 3F-tagged RPRDs in buffers containing increasing NaCl concentration. The bait proteins (red), as well as major interactors, are indicated with arrows. Protein marker sizes are shown on the left.

Taken together, RPRDs interactions with each other and with Pol II stay at the core of their binding properties.

### RPRD1B forms mutually exclusive complexes with RPRD1A or RPRD2

Our data indicate the existence of two separate complexes, RPRD1B-RPRD1A and RPRD1B-RPRD2, therefore we aimed to characterize these interactions in more detail. First, we explored whether there is a competition between RPRD1A and RPRD2 in binding to RPRD1B. For that, we performed IP of each RPRD1B, RPRD1A or RPRD2 upon the depletions of the two other RPRDs. SDS-PAGE analysis of precipitated fractions revealed that depletion of RPRD1A did not increase RPRD1B-RPRD2 interaction, neither did depletion of RPRD2 increase the formation of RPRD1B-RPRD1A complex (Figure 4A). Next, we used the RPRD1A-3F cell line to perform a two-step IP, in which tagged RPRD1A was precipitated first, with anti-FLAG antibody, and the flow-through fraction was used to precipitate RPRD2 using a commercial anti-RPRD2 antibody. In parallel, we performed an analogue experiment in which RPRD2 was precipitated first, followed by RPRD1A-3F precipitation from the flow-through fraction. Western blot analysis showed that the amount of co-immunoprecipitated RPRD1B either by RPRD1A-3F or RPRD2 was the same, independent of the fraction used (Figure 4B). Our experiments indicated that RPRD1A and RPRD2 do not compete with each other and there is a pool of RPRD1B available for RPRD1A and RPRD2 binding.

**Figure 4.**
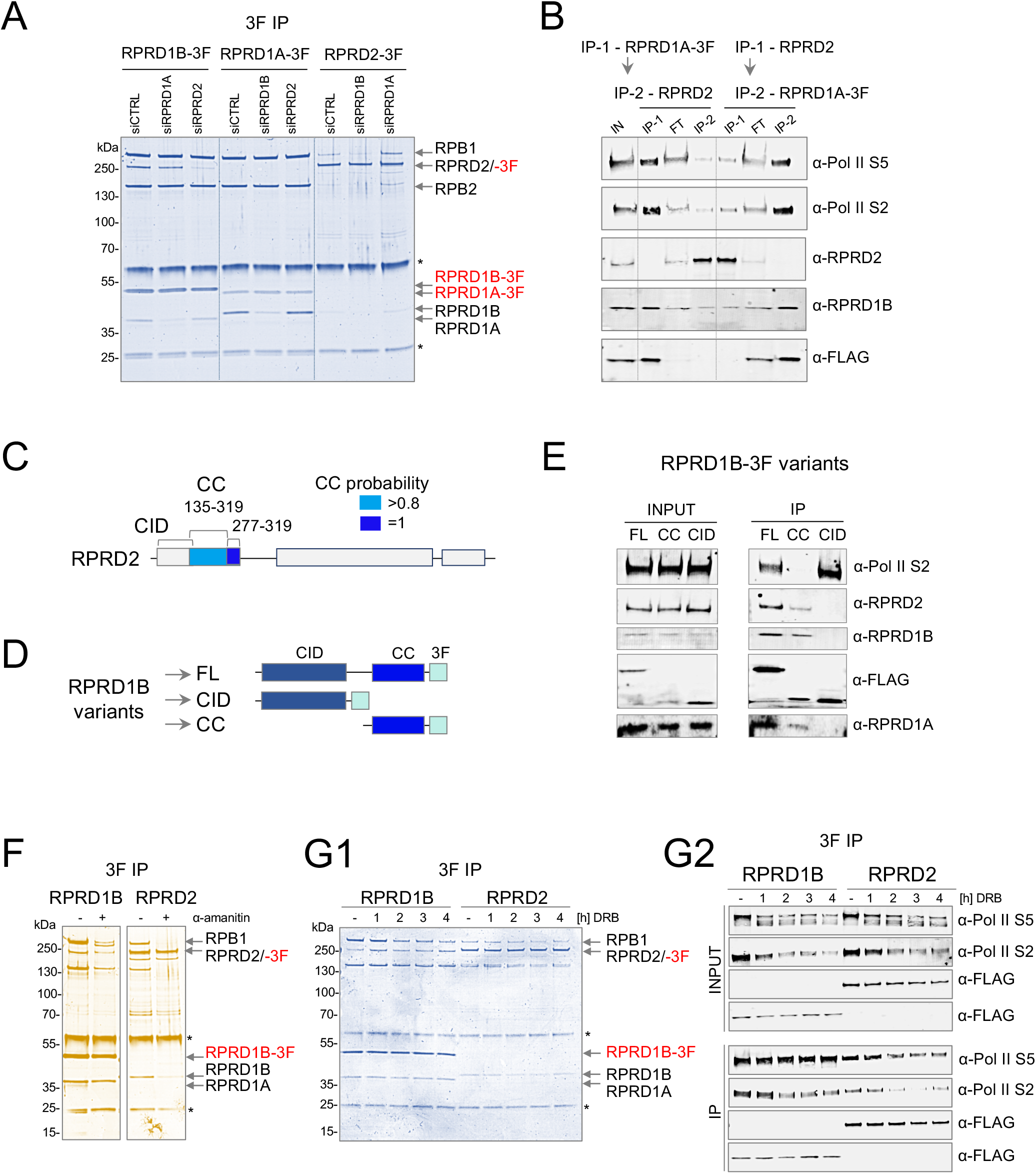
The interdependence of RPRD2, RPRD1B and RPRD1A interactions. **A.** A Coomassie stained SDS-PAGE gel analysis of proteins co-purified with 3F-tagged RPRDs upon depletions of other RPRDs. The bait proteins (red), as well as major interactors, are indicated with arrows. Protein marker sizes are shown on the left. Asterisk (*) denote heavy and light IgG contaminants. **B.** A western blot analysis of the indicated proteins immunoprecipitated with anti-FLAG and anti-RPRD2 antibodies from the RPRD1A-3F cell line. IP-1 – first IP from the input fraction; IP-2 – second IP from the flow-through fraction; IN – input fraction; FT – flow-through fraction **C.** A schematic representation of the domain structure of RPRD2 and the potential position of the CC domain based on the probability calculated by the Marcoil algorithm. CID – CTD-interacting domain; CC – coiled-coil domain**. D.** A schematic representation of the domain structure of the full-length (FL) RPRD1B-3F and its CID and CC domain mutants. CID – CTD-interacting domain; CC – coiled-coil domain**. E.** A western blot analysis of the indicated proteins co-purified with the RPRD1B-3F and its domain mutants. FL – full-length; CID – CTD-interacting domain; CC – coiled-coil domain. **F.** A silver stained SDS-PAGE gel analysis of proteins co-purified with 3F-tagged RPRD1B and RPRD2 after α-amanitin treatment for 24 hours. The bait proteins (red), as well as major interactors, are indicated with arrows. Protein marker sizes are shown on the left. Asterisk (*) denote heavy and light IgG contaminants. **G1.** A Coomassie stained SDS-PAGE gel analysis of proteins co-purified with 3F-tagged RPRD1B and RPRD2 upon treatment with DRB for the indicated time. The bait proteins (red), as well as major interactors, are indicated with arrows. Protein marker sizes are shown on the left. Asterisk (*) denote heavy and light IgG contaminants. **G2.** A western blot analysis of indicated proteins co-purified with 3F-tagged RPRD1B and RPRD2 upon treatment with DRB for the indicated time.

Next, we investigated how RPRD1B interacts with RPRD2. RPRD1B-RPRD1A heterodimer has been shown to form the interaction via their CC domains (Ni et al., 2014). Considering high sequence similarity between the RPRD2 N-terminus and RPRD1A and RPRD1B proteins (Figure S1A), we searched for a CC domain in RPRD2. Three CC-domain predicting software (Delorenzi and Speed, 2002; Gruber et al., 2005; McDonnell et al., 2006) revealed a putative CC domain in RPRD2, spanning from 135-319 aa, located next to the CID domain, as is seen for RPRD1A and RPRD1B (Figure 4C and S4A). Therefore, we tested whether the CC domain of RPRD1B is also necessary for the interaction with RPRD2. Using the Flp-In T-REx 293 system, we engineered cells expressing 3F-tagged RPRD1B mutants consisting only of the CC or CID domain (Figure 4D). Immunofluorescent staining showed that, single domains localised to the cytoplasm or both the cytoplasm and the nucleus, thus, we added a nuclear localization signal from SV40 large T antigen to restore their nuclear localization (Figure S4B). Next, we induced the expression of RPRD1B variants by adding Tet to the growth medium for 24 hours. The accumulation of the CC domain was much lower than that of the CID variant, indicating that the CID domain is necessary to stabilise the protein (Figure S4C). Therefore, in order to maintain similar levels of all RPRD1B variants, we induced the expression of FL and the CID domain with 2 ng/ml and the CC domain with 50 ng/ml Tet. Western blot analysis of fractions co-immunoprecipitated with anti-FLAG antibody revealed that the CC domain interacts with endogenous RPRD1B, RPRD1A and RPRD2 but not with S2 phosphorylated Pol II (Figure 4E). The CID domain precipitated only with Pol II, while the FL RPRD1B, interacted with all RPRDs and Pol II. Interestingly, the CID domain co-precipitated more Pol II than the FL protein, indicating that the presence of the CC domain and/or interaction with other RPRDs have an inhibitory effect on the binding to the Pol II. Similar patterns for the CC and CID domains’ binding properties was observed for RPRD1A variants. RPRD1A CC domain precipitated endogenous RPRD1A, RPRD1B and small amounts of Pol II, while RPRD1A CID domain precipitated Pol II only (Figure S4D).

Next, we tested if RPRD1B and RPRD2 require Pol II for interaction. For that, we reduced the levels of Pol II by treating cells for 24h with α-amanitin that directs specific Pol II degradation via the ubiquitination pathway (Lee et al., 2002). Next, we performed IP of either RPRD1B or RPRD2 and analysed the precipitated fractions on SDS-PAGE. The treatment altogether affected the formation of the RPRD complexes, as shown previously (Ni et al., 2014), possibly due to general deleterious effects caused by Pol II degradation. However, RPRD1B-RPRD2 and RPRD1B-RPRD1A interactions were sustained in the absence of Pol II, confirming that its presence is not required for the formation of RPRDs complexes (Figure 4F).

Next, we investigated if RPRD1B-RPRD2 interaction depends on active transcription. We blocked transcription elongation with DRB and tested RPRD1B and RPRD2 interactions by IP followed by SDS-PAGE and western blot analyses (Figure 4G1 and 4G2). After 4 hours of DRB treatment, RPRD1B co-purified less Pol II subunits when compared to time 0, while its interactions with RPRD1A and RPRD2 were unaffected. Consistently, a lack of active Pol II affected the RPRD2-Pol II interaction but not the RPRD1B-RPRD2 complex. This indicates that RPRD1B and RPRD2 bind to actively transcribing Pol II, however, this is not required for their interaction.

Finally, we tested RPRDs interactions in Flp-In T-REx 293 cells OE either RPRD2, RPRD1B or RPRD1A. We gradually increased the RPRDs OE using the increasing Tet concentration in the media. We then performed IP followed by SDS-PAGE analysis. RPRD2 OE did not affect the composition of its main complex but the quantity of each interacting partner increased proportionally to the level of the bait (Figure 5A). For RPRD1B, low OE (2 ng/ml Tet) led to a proportional increase in co-purification of its partners; however, high OE (50 ng/ml Tet) led to increased co-precipitation of Pol II but almost completely abolished the binding to RPRD1A and RPRD2 (Figure 5B1). Such a decrease in RPRDs interactions was not caused by saturation of endogenous pools of RPRD1A and RPRD2 as the two proteins remained in the flow-through fraction (Figure 5B2). We observed a similar effect when overexpressing RPRD1A, which resulted in a sustained interaction with Pol II but lower interaction with RPRD1B (Figure 5C1 and 5C2).

**Figure 5.**
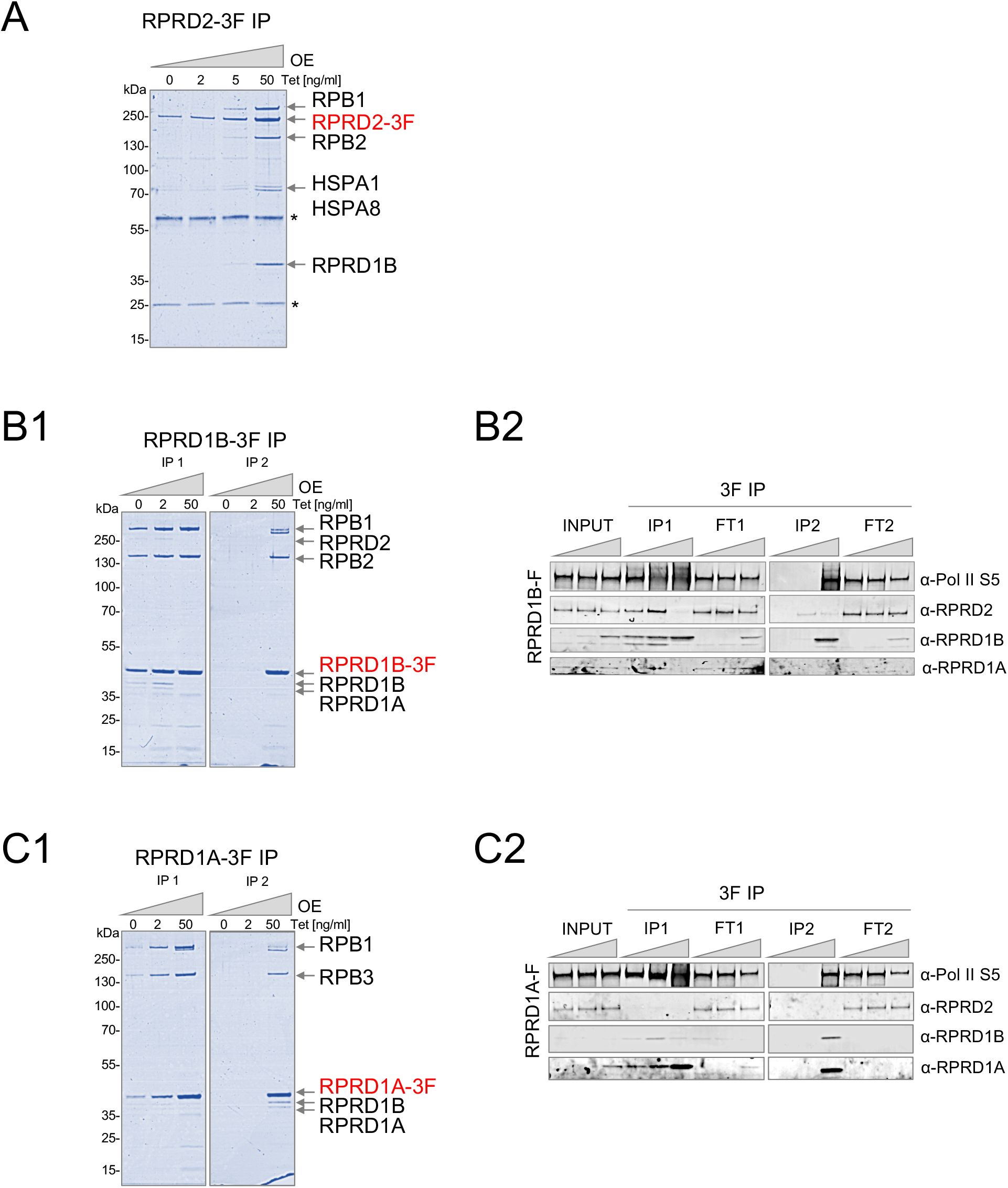
Differential formation of RPRD complexes enforced by overexpression. **A.** A Coomassie stained SDS-PAGE gel analysis of proteins co-purified with 3F-tagged RPRD2 upon its overexpression in the Flp-In T-REx cell line. Overexpression was induced for 24h with indicated concentrations of tetracycline. The bait protein (red), as well as major interactors, are indicated with arrows. Protein marker sizes are shown on the left. Asterisk (*) denote heavy and light IgG contaminants. **B1.** A Coomassie stained SDS-PAGE gel analysis of proteins co-purified with RPRD1B-3F upon its overexpression in the Flp-In T-REx cell line. Overexpression was induced for 24h with indicated concentrations of tetracycline. The second IP was performed on the flow-through fraction derived after the first IP. The bait protein (red), as well as major interactors, are indicated with arrows. Protein marker sizes are shown on the left. Asterisk (*) denote heavy and light IgG contaminants. **B2.** A western blot analysis of the experiment showed in B1. The input, IP and flow-through fractions were probed with the indicated antibodies. **C1.** As in B1 but for RPRD1A-3F. **C2.** As in B2 but for RPRD1A-3F.

### Biological consequences of RPRD2 depletion

The possible functions of RPRD1A and RPRD1B, but not RPRD2, have been studied in a wider biological context (Li et al., 2021). Thus, we sought to understand how RPRD2 affects cellular pathways. First, we tested the impact of RPRD2-dependent deregulation of RNA synthesis on general gene expression. The goal of this experiment was to gain insight into how the cell reacts to decreased RPRD2 levels by accordingly adjusting gene expression. We used the 3’RNA-seq data to search for differentially expressed genes resulting from RPRD2 depletion. We found that RPRD2 KD affected expression of 64 genes (Figure 6A). Interestingly, among the deregulated genes were *TOR3A* (alias ADIR, ATP-Dependant Interferon Response Protein 1), a protein involved in HIV-1 restriction (Luo et al., 2016); *APLN* (Apelin) that inhibits HIV-1 entry to the cell (Zou et al., 2000) and *CD70* that plays a role in the maintenance of T cell immunity (Izawa et al., 2017; Welten et al., 2013) (Figure 6B and Figure S6A). The 3’RNA-seq data from all three RPRD KDs revealed that the lower levels of *TOR3A* and *APLN* were particularly associated with RPRD2 but not with RPRD1A or RPRD1B depletions (Figure 6C). *CD70* was also significantly lowered only in RPRD2 KD (Figure S6B). Gene ontology (GO) terms analysis of genes that were significantly affected by RPRD2 KD (Table S3) revealed that apart from transcriptional pathways, the second most significant group of affected processes, were those involved in immunological responses (Figure 6D). We also analysed GO terms associated with proteins interacting with RPRD2 identified by our MS analysis (Figure 2B). The results consistently revealed that RPRD2 interactors are part of immune pathways responding to viral infections (Figure 6E). Other genes downregulated by RPRD2 KD are involved in the control of cell growth, for example in the regulation of apoptosis (*BAX*) (Kale et al., 2018) or associated with carcinogenesis ncRNA (*SNHG10*/*SCARNA13*) (Lan et al., 2019) (Figure 6F). This, and previous reports showing that other RPRDs stimulate or inhibit growth via opposite regulation of the cell cycle (Li et al., 2021), prompted us to investigate how RPRD2 levels drive changes in cellular growth. We combined RPRD2 KD with the synchronisation of the cells at the transition from G1 to S phase using double thymidine block (DTB) (Figure 6G). High concentrations of thymidine interrupt the deoxynucleotide metabolism pathways through competitive inhibition, thus blocking DNA replication. Cells released from the arrest and allowed to progress through the cell cycle were collected at four- and eight-hour time points and the DNA content was assessed by propidium iodide (PI) staining followed by flow cytometry (Figure 6F). On the first checkpoint, both the KD and control samples displayed a similar pattern in the progression to S phase. 8 hours post release, 40% of control cells were in G2/M phase compared to 65% of RPRD2 KD cells. Thus, we concluded that RPRD2 KD resulted in faster progression through S phase into G2/M. This highlights RPRD2 as a critical cell growth factor further underlining its role as a global transcription silencer involved in the regulation of gene expression.

**Figure 6.**
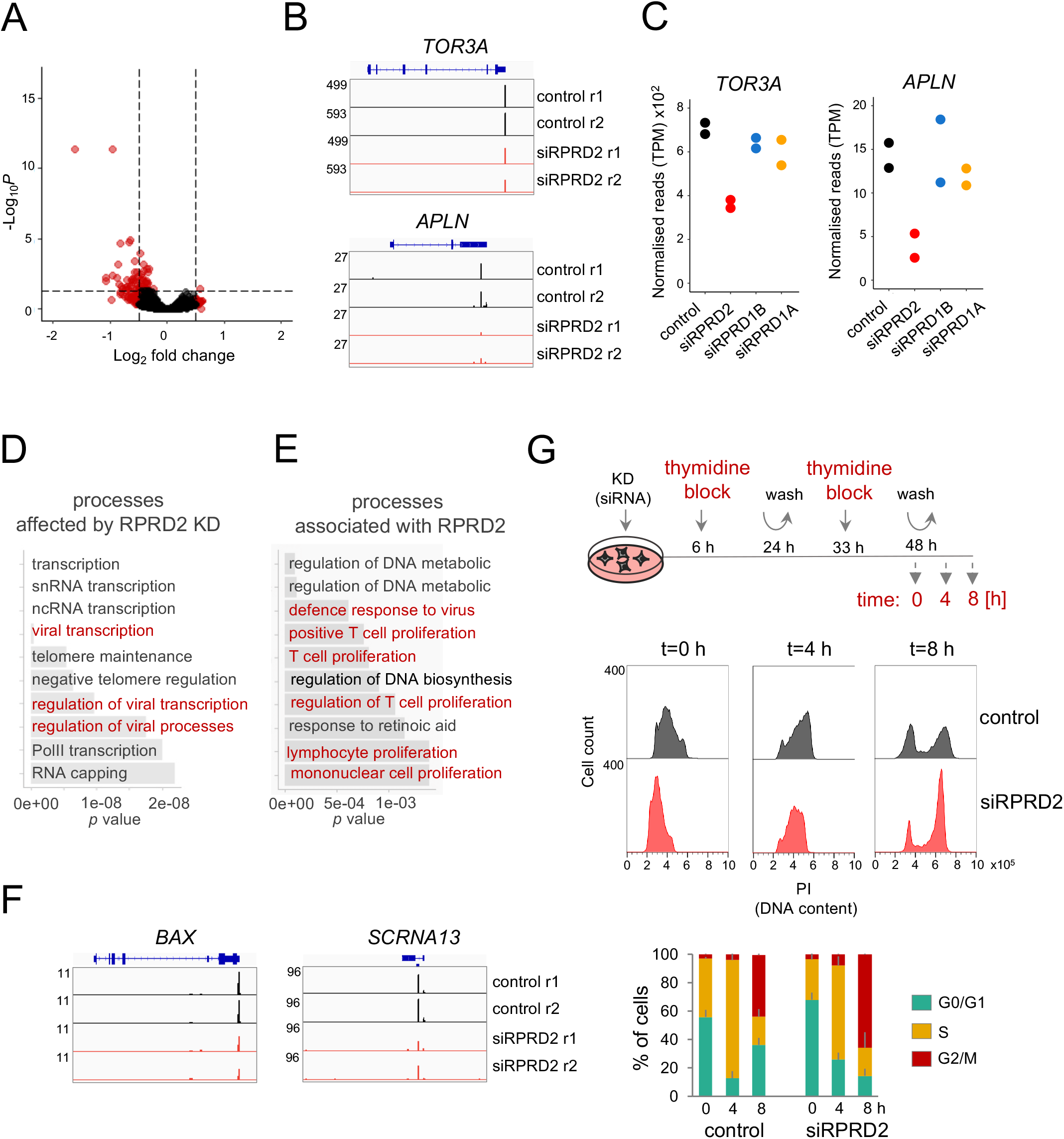
Biological roles for RPRD2. **A.** Differentially expressed genes in RDRD2 KD in relation to control. Red denotes genes with adjusted p-value < 0.05 or *log_2_* fold change value < −0.5 or > 0.5. **B.** IGV (Integrative Genomics Viewer) tracks showing the levels of anti-viral factors, *TOR3A* and *APLN,* mRNA in control and RPRD2 KD (each with two replicates). **C.** A dot plot showing the 3’RNA-seq analysis of the *TOR3A* and *APLN* mRNA levels in control and RPRD KDs. For each condition, two dots of the same colour represent two replicates. **D and E.** Gene ontology (GO) terms associated with (D) genes differentially expressed in RPRD2 KD identified in 3’RNA-seq, and (E) with RPRD2 protein interactors identified by mass spectrometry. In red are denoted terms related to the cellular anti-viral processes. **F.** IGV tracks showing the expression levels of the apoptotic factor *BAX* and cancer-related *SCRNA13*, identified as differentially expressed in RPRD2 KD 3’RNA-seq (each with two replicates). **G.** The analysis of the cell cycle progression in RPRD2 KD. *Top* The diagram showing the experimental approach. Control and siRNA-treated cells were subjected to a double thymidine block (DTB) to synchronise the cell cycle in the G1/ S phase. Samples were collected after the release from DTB at the indicated time points. *Middle* Histogram of cells stained with propidium iodide (PI) showing DNA content distribution at indicated time points after the release from DTB in control and RPRD2 KD. *Bottom* A bar plot showing the percentage of cells in each cell cycle phase at indicated times after the DTB release in control and RPRD2 KD. Error bars represent the standard deviation calculated from two independent experiments.

## DISCUSSION

The expression of many genes is beneficial in only particular conditions and for most of the time they may have deleterious effects on a living cell if expressed in excess. The option to downregulate transcription allows maintaining gene expression relatively low but not completely shut down. This, in turn, offers the possibility to robustly turn on gene expression when necessary (Adelman et al., 2009). Gene expression can be downregulated at the transcriptional level by many factors. The simplest way is to prematurely terminate transcription, however to restore transcription, all early steps including the formation of the preinitiation complex, and transcription initiation processes have to be repeated (Shandilya and Roberts, 2012). Pol II pausing or maintaining low level of transcription is more versatile and allow for a rapid shift from the “low” to “high” expression and *vice versa* (Adelman and Lis, 2012; Core and Adelman, 2019; Danko et al., 2013; Gilchrist et al., 2012). Such transitions make up the foundations of general survival strategies, most vividly underlined during responses to stresses like heat, oxidants or viral infections (de Nadal et al., 2011; Katze et al., 2002).

We found that Pol II CTD interactors, RPRD proteins, RPRD2 and RPRD1B in particular, act as negative transcription factors that affect the levels of newly transcribed and nascent RNAs. RPRDs depletion or overexpression exerted opposite impacts on RNA synthesis. Low levels of RPRD increased the accumulation of nnsRNAs, while overexpression of RPRD2 and RPRD1B, and to lesser extent RPRD1A, suppressed RNA synthesis. We observed the most profound effect on transcription of both KD and OE in the case of RPRD2. This protein is larger than RPRD1A and RPRD1B and contains Ser- and Pro-rich domains that may be used for the recruitment of other transcriptional regulators. The effects of the RPRD1A KD on nnsRNA were moderate and in the case of OE experiments, opposite to RPRD2 and RPRD1B OE. Interestingly, RPRD1B impact on the transcriptome was intermediate when compared to RPRD2 and RPRD1A KD or OE. The explanation of this may arise from our proteomics studies. We found that these three proteins can form two independent complexes. RPRD1B interacts with either RPRD1A or RPRD2 via its CC domain, but an interaction between RPRD1A and RPRD2 was not detected in our experiments. The RPRD1B-RPRD1A complex has been reported to recruit RPAP2 phosphatase that dephosphorylates Pol II CTD at S5 in the early stages of transcription and facilitate the transition into elongation (Ni et al., 2014). The fact that RPRDs form interchangeable complexes while also may function separately as monomers, adds to the complexity of how these proteins regulate transcription. We found that RPAP2 interacts with RPRD1B and RPRD1A but not with RPRD2. Consistently, only RPRD2 but not RPRD1B and RPRD1A KD increased transcription of snRNAs which require RPAP2 for their synthesis (Egloff et al., 2012). Thus, we do not rule out that RPRD1B and RPRD1A, in complex, may act as positive transcription regulators. RPRD1A KD resulted in decreased RPAP2 interaction with Pol II and association with selected promoters (Ni et al., 2014). The investigation into if RPAP2 only requires RPRD1A or RPRD1B-RPRD1A heterodimer, will be crucial to understanding the roles of the RPRDs in the promotion of transcription.

We show that two mutually exclusive complexes, RPRD1B-RPRD1A and RPRD1B-RPRD2, co-exist in the cell. Their interactions are mediated by the CC domains and are Pol II – independent. However, when overexpressed, RPRD1B and RPRD1A preferentially bind to Pol II, losing interactions with other partners. Overexpression of RPRD2, on the other hand, increases the level of all its interactors. Such disparity indicates different mechanisms of Pol II-RPRDs connections and may be the cause of certain differences observed in the transcriptional regulation by the three proteins. Thus, it is essential that cells maintain the levels of RPRD proteins balanced. Overexpression of RPRD2 may actually increase the presence of the RPRD1B-RPRD2 dimer on Pol II and therefore, outcompete RPRD1A-RPRD1B. In the case of RPRD1B OE, it seems that ‘free’ RPRD1B preferentially interacted with Pol II, without its usual partners, RPRD1A or RPRD2. In both scenarios, transcription is severely affected.

How can RPRD1B and RPRD2 downregulate transcription? The only protein that co-purified with all three RPRDs, apart from RPB1 and RPB2, was Pol II subunit POLR2M, also called GDOWN1. This protein is present on a number of promoters and binds directly to Pol II RPB1, RPB2, RPB3, and RPB10 subunits. Cryo-EM analysis showed that GDOWN1 overlaps with the TFIIB and TFIIF interaction sites on RPB1 and RPB2 (Jishage et al., 2018). Notably, RPB1 and RPB2 are strongly associated with RPRD proteins (Figure 3D and S3B). Similar to our observations, GDOWN1 KD causes an increase in transcription (Cheng et al., 2012; Jishage et al., 2012). Moreover, in *Drosophila*, GDOWN1 is inversely co-related with transcriptional activation in early embryos (Jishage et al., 2018). It is possible that the RPRD proteins may regulate GDOWN1 recruitment to Pol II and so transcription rates. RPRDs were also reported to interact with the PAF1 complex however this interaction may require additional factors as the subunits were not detected in stringent purification conditions. PAF1 has been shown to both stimulate and inhibit transcription (Francette et al., 2021), thus, it may be a suitable partner for transcription stimulation and repression performed by interchanging RPRD complexes. Thus, RPRDs may associate with various factors to adjust RNA synthesis levels in a context dependent manner.

Our findings indicate that RPRD2 may act as an efficient transcriptional regulator and a negative growth factor. Overexpression of RPRD2 led to the inhibition of transcription and affected cellular growth. It is possible that RPRD2 can act independently of RPRD1B, however, due to its deleterious activity, it may require a factor, like RPRD1B, to modulate its activity or recruitment to the transcribing complex. We found that RPRD2 is bound by two heat shock proteins (HSP), however, further investigations are needed to understand the function of this interaction. HSPs have been reported to perform various functions ranging from protein stabilisation to directing them to degradation (Fernandez-Fernandez et al., 2017). Therefore, HSPs may regulate RPRD2 levels and so its activity in the cell. RPRD2 anti-transcriptional activity may also be effectively used by cellular defence systems against the expression of exogenous DNA. Previous reports showed that RPRD2 (called REAF) displays antiviral properties against HIV-1. RPRD2 can be quickly displaced to the cytoplasm and supress reverse transcription of viral RNA into DNA (Gibbons et al., 2020). We revealed that RPRD2 antiviral activities may be expanded on the broad control of cellular pathways. For example, one of the genes whose expression was significantly affected by RPRD2 was *TOR3A*, interferon-induced chaperone-like ATPase (Dron et al., 2002), a protein shown to compromise the infectivity of HIV-1 (Luo et al., 2016). In general, RPRD2 depletion affected the expression of a number of genes involved in immunological responses. We speculate that RPRD2 may act generally during stress response and, thus, regulate the expression of genes that are induced by cellular stress, including viral infections.

The lack of control in RPRDs cellular levels has been suggested to drive carcinogenesis implying their important roles in regulation of gene expression (Li et al., 2021; Lu et al., 2012; Wu et al., 2010). However, deregulation of nascent transcription in HEK293T cells was not fully mirrored by steady-state RNA levels. Insufficient or excessive RNA synthesis may be buffered by RNA stability pathways (Haimovich et al., 2013; Sun et al., 2013). Although the mRNA levels are adjusted, cells with faulty transcriptional control may not react efficiently to changing metabolic conditions, cellular signals or stress. Thus, the errors in gene expression may accumulate over time and reach a point when a cell loses control of gene expression, starts proliferating uncontrollably, escapes apoptosis and becomes a cancer cell.

Overall, we report a new function for RPRD1B, RPRD1A and RPRD2 proteins in the control of RNA synthesis. Our results show that RPRD2 and RPRD1B, and to lesser extent RPRD1A, act as negative transcription factors, adding to complexity of transcription regulation.

## ACKNWOLEDGMENTS

PG lab (KW, HC, MS, HF) was supported by a Sir Henry Dale Fellowship from the Wellcome Trust and the Royal Society (200473/Z/16/Z). HC was supported by a studentship from the Ministry of National Education, Republic of Turkey. DH is funded by EPSRC (grant EP/T002794/1), BBSRC (grants BB/L006340/1 and BB/M017982/1); MJ is supported by Midlands Doctoral Training Partnership for a studentship to M.J. funded by BBSRC BB/M01116X/1. We thank Charlotte Lal for helping with cell line construction, Anshita Goel and Genomics Birmingham at the University of Birmingham for assistance with NGS, and the Advanced Mass Spectrometry Facility at the University of Birmingham for assistance with MS.

## CONTRIBUTION

K.W. and H.C. performed most experiments in this study. K.W. performed proteomics studies, cell line construction and functional assays. H.C. performed NGS experiments and functional assays. M.S., M.J. and D.H. performed bioinformatics analyses. H.F. constructed the cell lines. R.A. performed initial analyses. J.W. and M.Sa. supported FASC experiments. M.S. and H.F. participated in writing of the manuscript. K.W. and P.G. designed the experiments and wrote the manuscript.

## RESOURCE AVAILABILITY

Lead contact: Further information and requests for resources and reagents should be directed to the lead contact, Pawel Grzechnik (p.l.grzechnik(at)bham.ac.uk).

Data and code availability: The dataset generated during this study are available at GEO.

Accession numbers: 4sU-seq: GSE178213; 3’seq: GSE178495.

---

**Figure S1.**
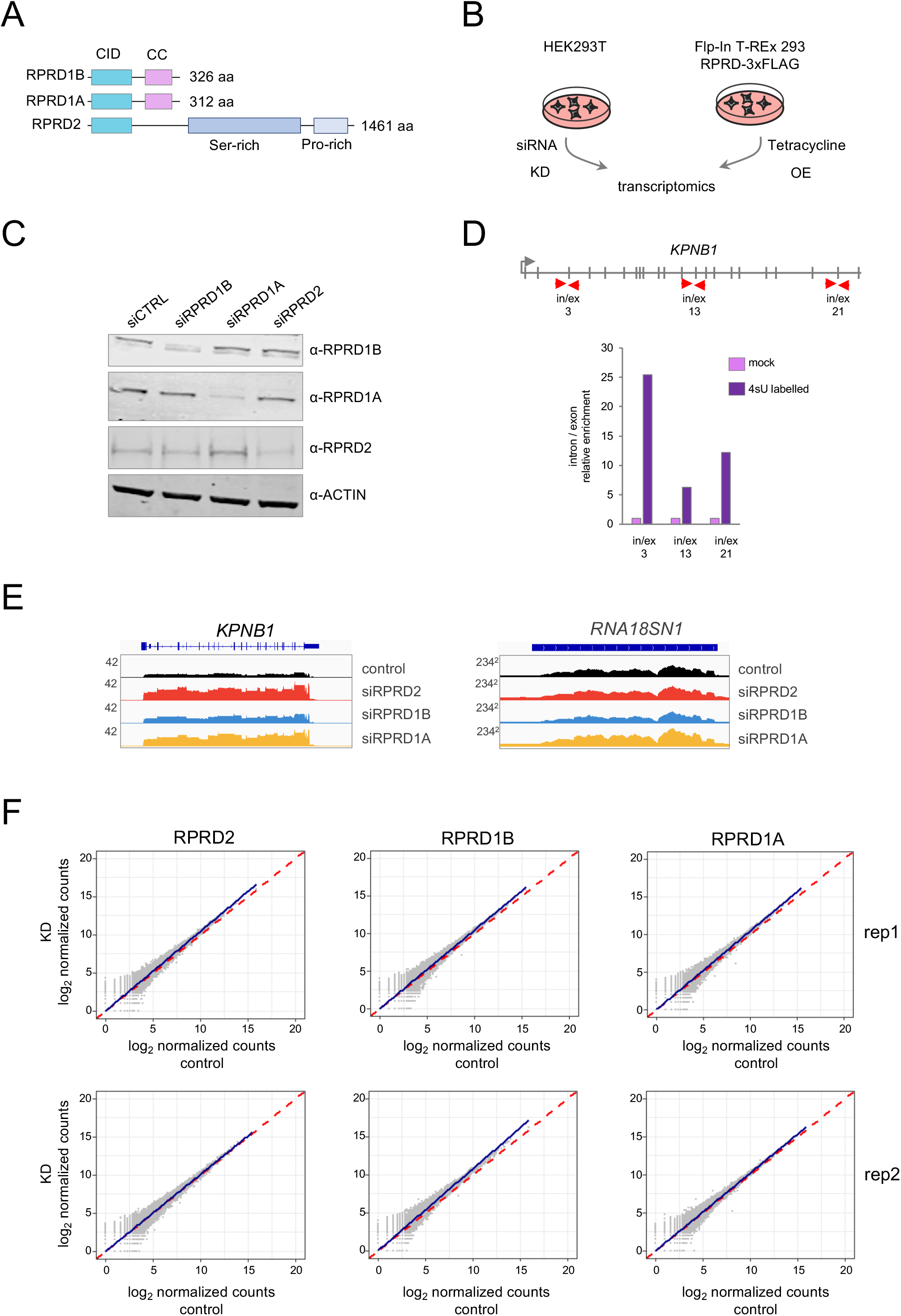
Related to Figure 1. **A.** Schematic representation of the size and domain structure of RPRD proteins. CID – CTD-interacting domain; CC – coiled-coil domain; Ser-rich – serine-rich domain; Pro-rich – proline-rich domain**. B.** Experimental approach to study the impact of depletion and overexpression of RPRD proteins on transcription regulation. To deplete RPRDs, HEK293T cells were treated with siRNA. To overexpress RPRDs, Flp-In T-REx 293 system was used for stable integration of RPRD genes copies under the control of tetracycline-inducible promoter. KD – knock-down; OE – overexpression. **C.** Western blot analysis of the indicated proteins in HEK293T cells treated with control and siRNA against RPRD1B, RPRD1A and RPRD2. **D.** *Top* Nascent and newly synthesised RNA is enriched during 4sU RNA labelling and precipitation; RT-qPCR analysis. A schematic representation showing introns and exons of the *KPNB1* gene. Red arrows point to the positions of the amplicons spanning over intron / exon junctions, tested in the analysis. *Bottom* Bar plot showing relative accumulation of *KPNB1* pre-mRNA in the mock (pink) and 4sU labelled (purple) samples. **E.** IGV (Integrative Genomics Viewer) tracks showing the accumulation of nnsRNA levels for *KPNB1* mRNA (left) and *RNA18SN1* rRNA (right) in KD samples and the control (counts x10^5^). **F.** Levels of nnsRNAs upon RPRDs KD, 3’-seq analysis. Scatter plot presenting the values of *log_2_* normalised counts for each gene in the KD samples (y-axis) against the control samples (x-axis). The blue lines represent the regression trend. The dashed red diagonals denote 0 *log_2_* fold change. Data is shown for two replicates.

**Figure S2.**
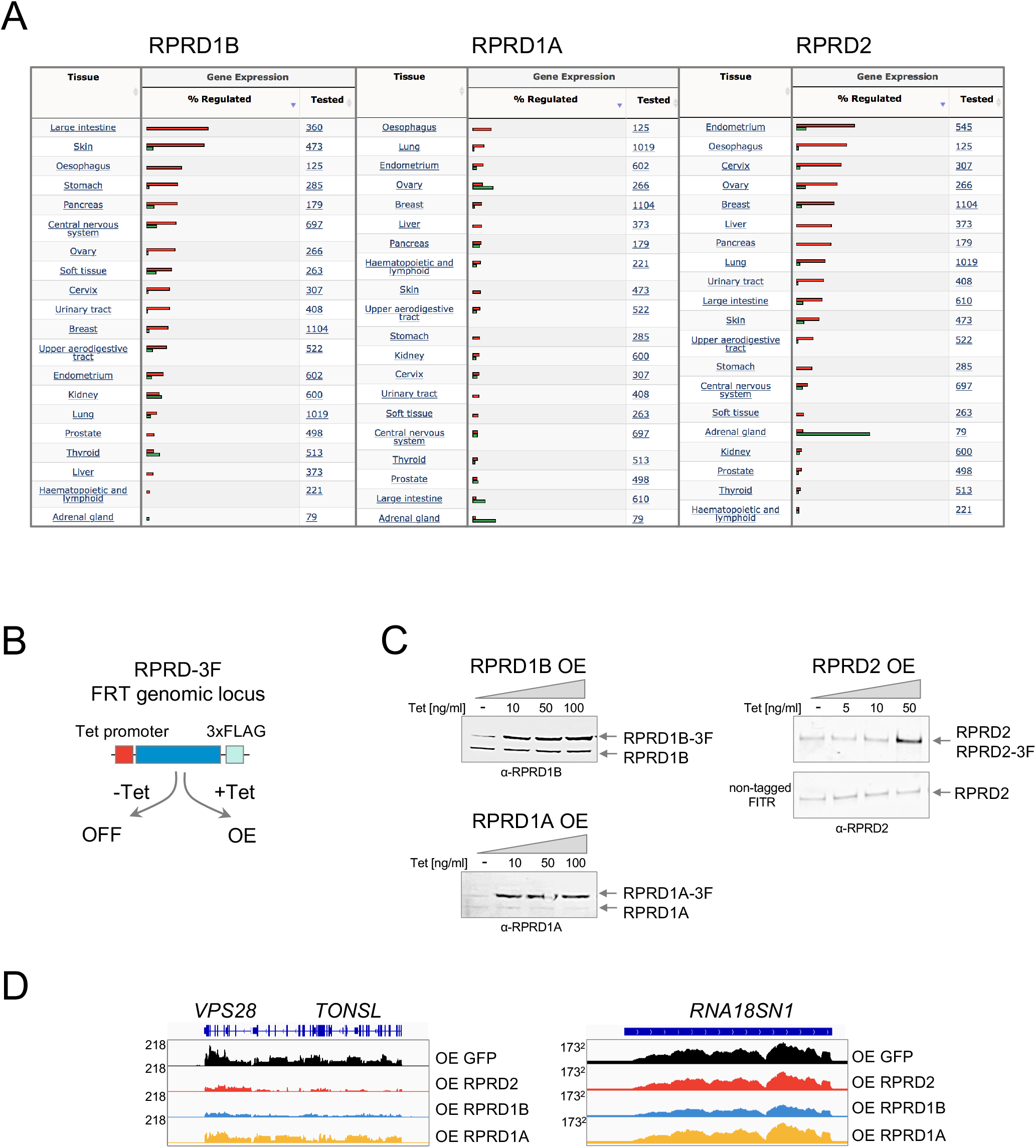
Related to Figure 2. **A.** RPRD1B and RPRD2 are often upregulated in cancer. The tables showing the distribution of RPRD proteins mutations across the primary tissue types, curated by Catalogue of Somatic Mutations in Cancer (COSMIC; www.cancer.sanger.ac.uk). Bars show the percentage of cancer tissue samples with deregulated RPRDs expression; red – upregulated, green – downregulated. **B.** A diagram showing the principle of overexpression of RPRDs using the Flp-In T-REx 293 (FITR) system. RPRDs coding sequences with added C-terminal 3F tag were integrated into FRT genomic locus under the control of tetracycline-inducible promoter. Addition of tetracycline to the growth media activates the gene expression. **C.** Western blot analysis of the indicated proteins after induction of overexpression with tetracycline. 3F-tagged and endogenous copies of RPRDs are indicated with arrows. In the case of RPRD2, the level of 3F-tagged protein overexpression is compared to the endogenous RPRD2 in the non-tagged FITR cell line. **D.** IGV (Integrative Genomics Viewer) tracks showing the accumulation of nnsRNA levels for *VPS28* and *TONSL* mRNA (left) and *RNA18SN1* rRNA (right) in OE samples and the GFP control (counts x10^5^).

**Figure S3.**
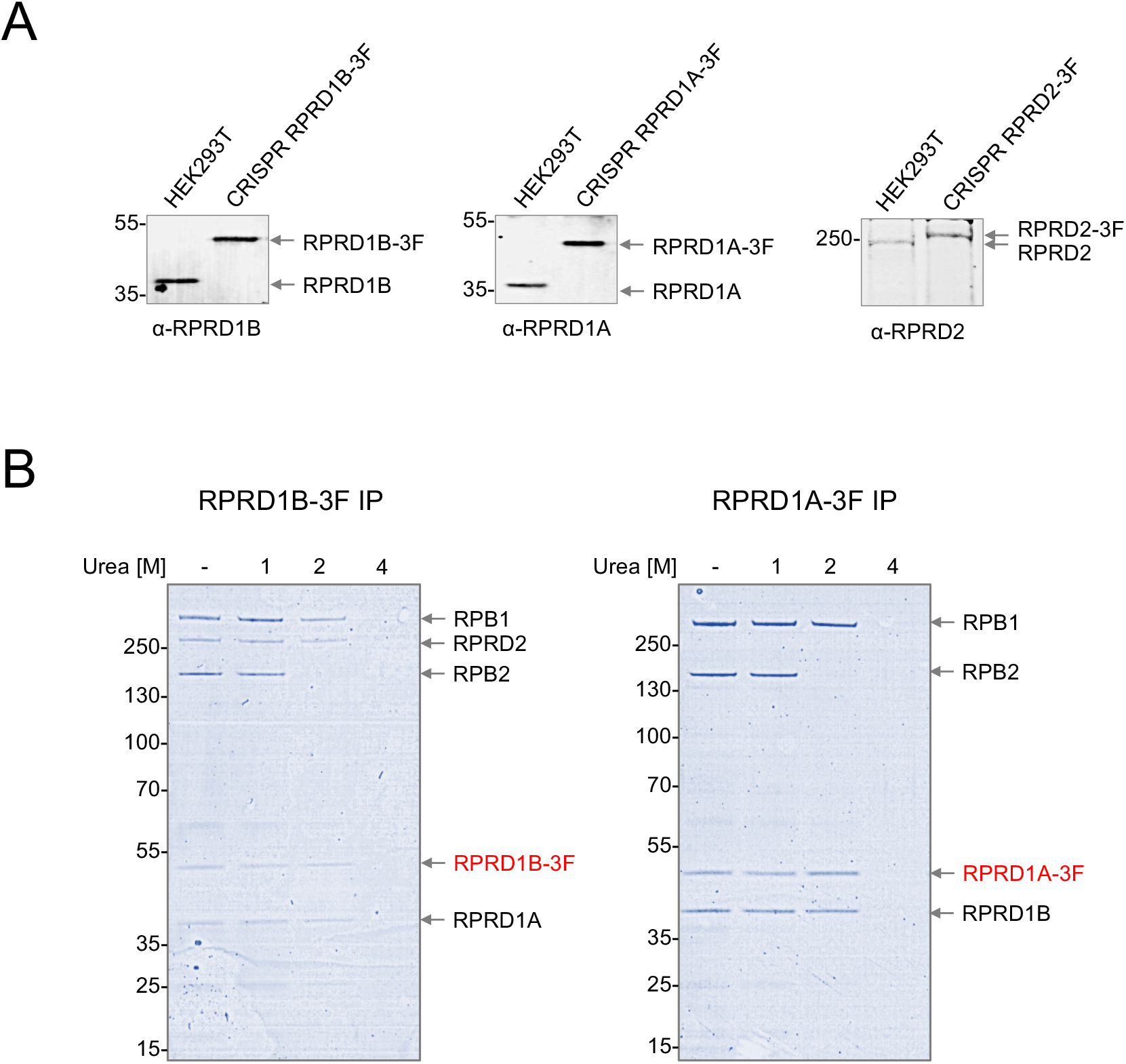
Related to Figure 3. **A.** Western blot analysis of the indicated RPRD proteins in 3F-tagged RPRD CRISPR cell lines in comparison to non-tagged HEK293T. 3F-tagged and endogenous copies of RPRDs are indicated with arrows. Protein marker sizes are shown on the left. **B.** A Coomassie stained SDS-PAGE gel analysis of proteins co-purified with 3F-tagged RPRD1A and RPRD1B in buffers containing an increasing concentration of urea. The bait proteins (red), as well as major interactors, are indicated with arrows. Protein marker sizes are shown on the left.

**Figure S4.**
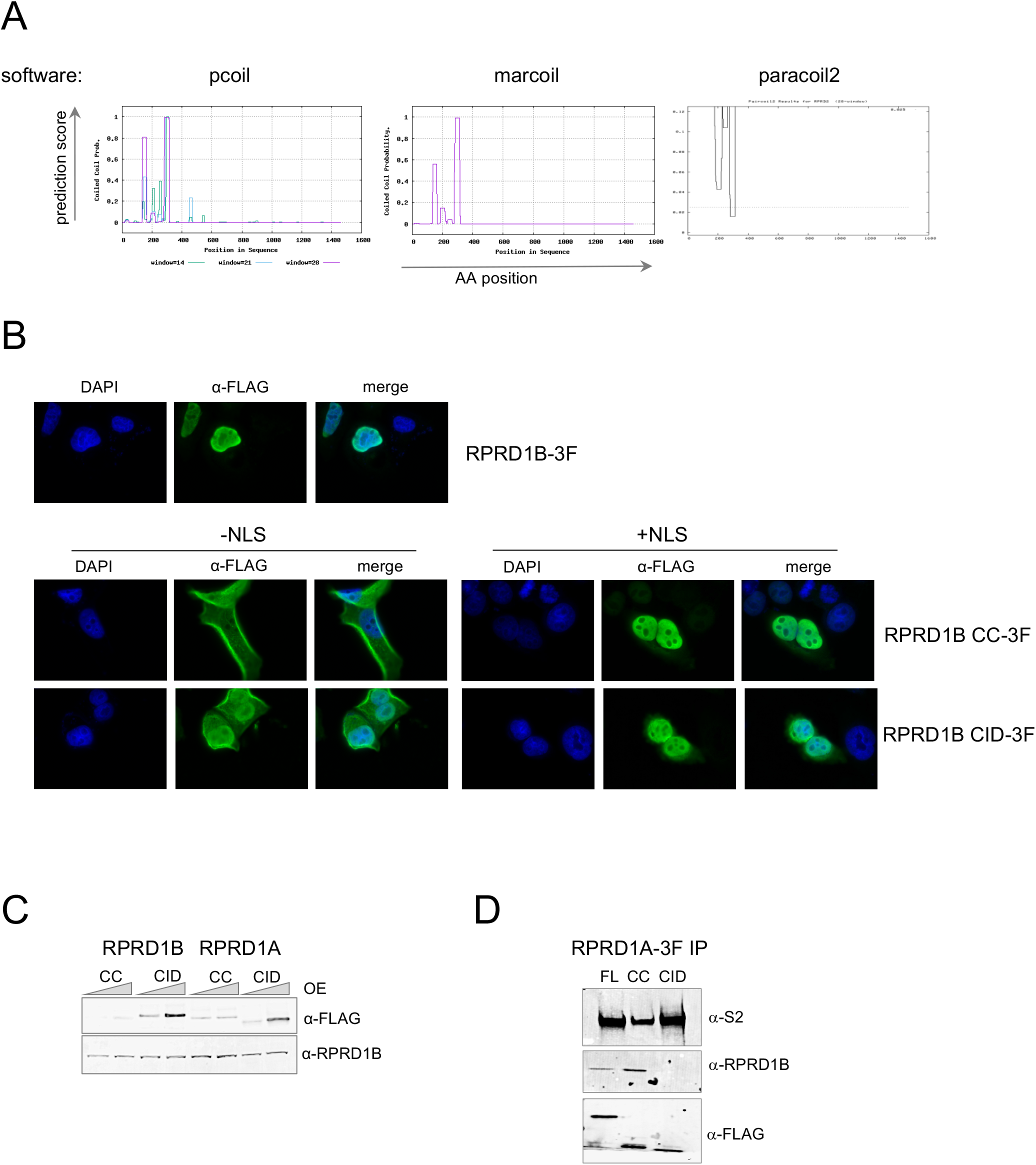
Related to Figure 4. **A.** Screenshots of the results of the coiled-coil (CC) domain predictions in RPRD2 using three different algorithms – pcoil, marcoil and paracoil2. All software model the CC presence in the same region within the 150-350 aa window. **B.** α-FLAG immunofluorescence microscopy analysis of HEK293T cells transiently expressing RPRD1B-3F (top) and RPRD1B CC and CID variants (bottom) with and without nuclear localisation signal (NLS) as indicated. 4 ,6-diamidino-2-phenylindole (DAPI) stain was included as a nuclear marker. **C.** Western blot analysis of the tetracycline-induced expression of 3F-tagged CC and CID RPRD variants in comparison to the levels of endogenous RPRD1B. **D.** Western blot analysis of the indicated proteins co-purified with the RPRD1A-3F and its domain mutants. FL – full-length; CID – CTD-interacting domain; CC – coiled-coil domain.

**Figure S6.**
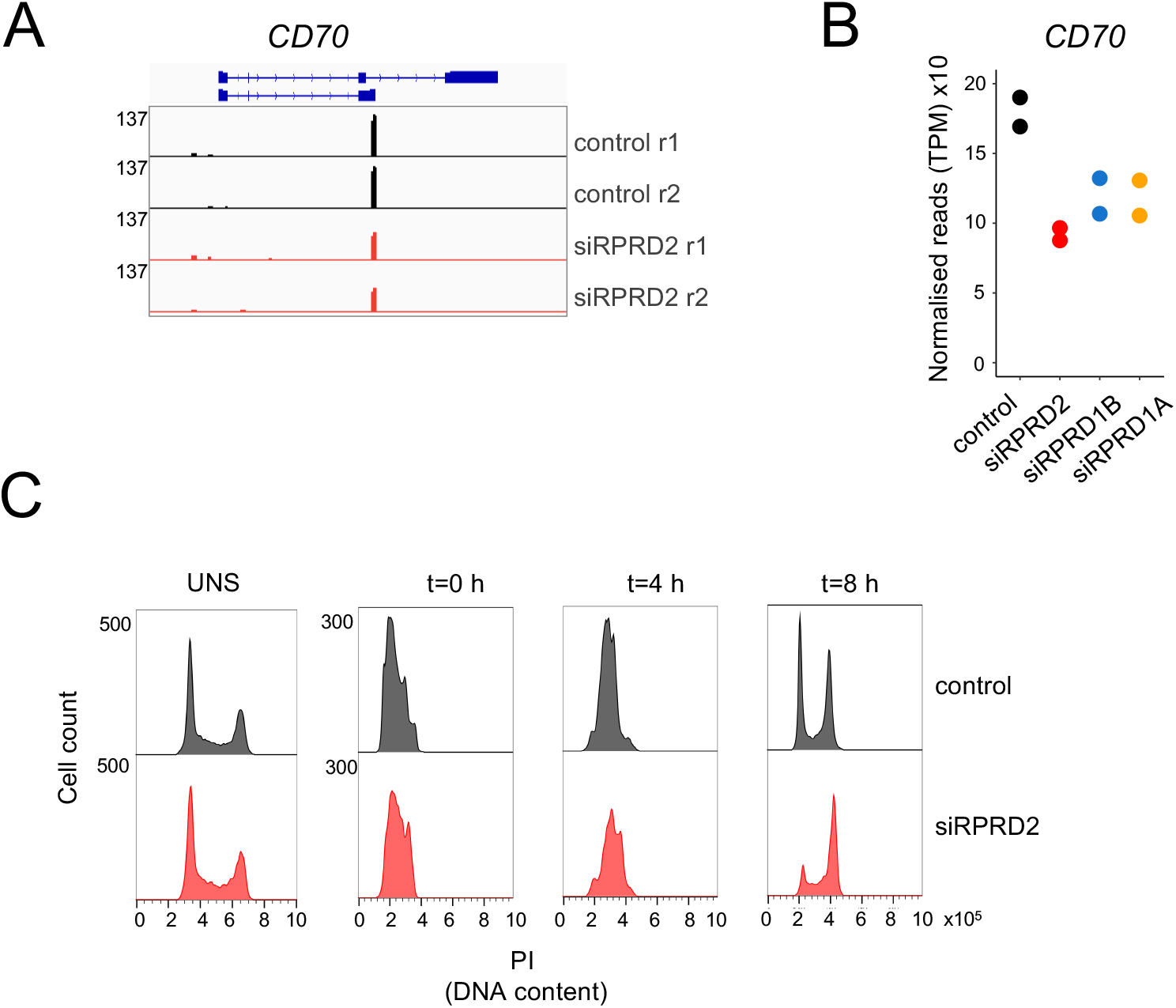
Related to Figure 6. **A.** IGV (Integrative Genomics Viewer) tracks showing the expression levels of *CD70* gene, identified as differentially expressed in RPRD2 KD 3’RNA-seq (two replicates). **B.** A dot plot showing the 3’RNA-seq analysis of the *CD70* mRNA levels in control and RPRD KDs. For each condition, two dots of the same colour represent two replicates. **C.** Histogram of cells stained with propidium iodide showing DNA content distribution in unsynchronised and G1/S synchronised cells (t=0 h) and at 4- and 8-hour time points after the release from synchronisation in control and RPRD2 KD. TRACK SHOWS

## METHODS

### Cell lines and cell culture

HEK293T and Flp-In T-REx 293 cells were maintained in 37°C and 5% CO2 in Dulbecco’s Modified Eagle Medium (Thermo Fisher, 41966029) supplemented with 10% Fetal Bovine Serum (Thermo Fisher, 10500064) and 1% Penicillin-Streptomycin (Merck, P0781).

### Creating 3F-tagged RPRD cell lines with CRISPR-Cas9

Three plasmids were prepared for tagging each RPRD: two donor plasmids bearing the 3F sequence targeting two alleles and a gRNA plasmid expressing the gRNA directing the Cas9 nuclease to the targeted loci. To make the gRNA plasmid, a gRNA sequence specific to the gene being tagged, was cloned into px330 vector using BbsI restriction site and subsequent ligation with T4 DNA ligase. To make donor plasmids left and right homology arms were amplified from genomic DNA and cloned into TOPO-TA/3F/blast and TOPO-TA/3F/hygro vectors using KpnI and NheI restriction sites for the left and MluI and XbaI restriction sites for the right homology arms, and subsequent ligation with T4 DNA ligase. Correct DNA sequence of final constructs was confirmed by Sanger sequencing. HEK293T cells were co-transfected with two donor and a gRNA plasmid using Lipofectamine2000 (Thermo Fisher, 11668019). 24h after transfection, selection media were added to the cells, supplemented with 100 μg/ml hygromycin (Thermo Fisher, 10453982) and 10 μg/ml blasticidin (Thermo Fisher, A1113902). After ~10 days single colonies appeared on the plate, which were then moved to separate dishes and further propagated under selection conditions. The presence of the tagged proteins was confirmed by western blot analysis.

### Creating 3F-tagged RPRD overexpression Flp-In T-REx 293 cell lines

Sequences of full-length and truncated RPRDs were amplified by PCR from cDNA and cloned into pcDNA5/FRT/TO vector containing C-terminal 3F-tag sequence using HindIII and EcoRV restriction sites and subsequent ligation with T4 DNA ligase. Correct DNA sequence of final constructs was confirmed by Sanger sequencing. Primers sequences are available upon request.

Flp-In T-REx 293 cells were co-transfected with pcDNA5/FRT/TO and pOG44 plasmids using Lipofectamine2000 (Thermo Fisher, 11668019). 24h after transfection, selection media were added to the cells, supplemented with 100 μg/ml hygromycin (Thermo Fisher, 10453982) and 10 μg/ml blasticidin (Thermo Fisher, A1113902). After ~10 days single colonies appeared on the plate, which were then further propagated under selection conditions. The presence of the tagged proteins was confirmed by western blot analysis.

### Immunofluorescent microscopy

Cells grown on cover slips fixed in 4% para-formaldehyde and permeabilized in 3% FBS, 0.5% Triton X-100 in PBS. Cover slips were then incubated with α-FLAG antibody (Merck, F7425) for 1h followed by incubation with goat α-rabbit Alexa Flour 488 (Thermo Fisher, A-11008) for 1h. Cover slips were mounted on glass slides with FluoroShield with DAPI (Merck, F6057).

### RNAi

Transfections were performed with siLentFect lipid reagent (Bio-Rad, 1703362) according to the manufacturer’s instructions, with siRNAs at a final concentration of 20 nM. siRNAs used were ON-TARGETplus SMARTpool for: RPRD1B (Horizon, L-013787-01-0020), RPRD1A (Horizon, L-007734-00-0020), RPRD2 (Horizon, L-021712-01-0020) and non-targeting control (Horizon, D-001810-10-50).

### Inhibition of Pol II activity

Cells were treated with 100 μM DRB (5,6-Dichlorobenzimidazole 1-β-D-ribofuranoside; Merck, D1916) or with 5 μg/ml α-amanitin (Merck, A2263) as indicated in the text.

### Colony forming unit assay

RPRD2-3F and GFP Flp-In T-REx cells were seeded in three 35-mm dishes each, at the density 100 cells/dish, and grown for 11 days in DMEM/FBS/PS containing tetracycline at indicated concentrations. Medium was changed every 3 days. After 11 days cells were fixed with 1% formaldehyde (Merck, F8775) for 15 min at room temperature with gentle rocking, and stained with 0.1% crystal violet stain (Merck, C0775).

### RNA extraction

Total RNA was extracted from cells using TRIzol reagent (Thermo Fisher, 15596026) according to the manufacturer’s protocol. Briefly, cells from one 10-cm plate were resuspended in 1ml TRIzol and incubated at room temperature for 5 min.

Next, 200 μl of chloroform was added and the mixture was vortexed for 15 s followed by 2-3 min incubation at room temperature. The sample was centrifuged for 15 min at 12000 x g at 4°C. RNA was precipitated with isopropanol for 10 min at room temperature. Next, RNA was pelleted by centrifugation for 10 min at 12000 x g at 4°C and washed once with 75% ethanol. RNA was dissolved in 50 μl nuclease-free water and the concentration measured with microvolume spectrophotometer. QuantSeq 3’mRNA-Seq Library Prep Kit REV for Illumina was used to prepare libraries for sequencing.

### RT-qPCR

Reverse transcription was performed with Super Script III Reverse Transcriptase (Thermo Fisher, 18080044) according to the manufacturer’s instructions. Random hexamers and anchored oligo-dT primers were used for cDNA synthesis. qPCR data were processed with the ddCt method, with normalization to both *GAPDH* mRNA levels and siRNA control sample. Primers sequences are available upon request.

### 4sU RNA labelling and precipitation

Cells were treated with 500 μM 4-thiouridine for 5 minutes before the RNA extraction was performed with TRIzol according to the manufacturers’ instructions. Total RNA was spiked-in with 4-thiouracil (4tU, Sigma, 440736)-labelled *S. cerevisiae* RNA (10:1 ratio). Biotinylation of the 4sU/4tU labelled RNA was performed with EZ-Link HPDP-Biotin (Thermo Fisher) for 30 minutes at 65°C. Biotinylated RNA was precipitated with chloroform/isopropanol and incubated with Dynabeads M-270 Streptavidin (Thermo Fisher) for 1 hour at room temperature. Beads were washed five times with Binding & Washing buffer (5 mM Tris-HCl (pH 7.5), 0,5 mM EDTA, 1 M NaCl) and 4sU labelled-RNA was eluted with 100 μM DTT, precipitated with glycogen and resuspended DNase/RNase free water. Enrichment of nascent RNA was confirmed with qPCR using exon–intron junction primers of *KPNB1* gene. Primers sequences are available upon request. NEBNext Ultra8482 II RNA Library Prep Kit for Illumina was used to prepare the sequencing libraries.

### RNA sequencing

The RNA sequencing was performed at the Genomics Facility at the University of Birmingham. Briefly, the concentrations and the average library size were determined using TapeStation (Agilent). The libraries were then diluted to 1.6 pM, and 1% of 20 pM PhiX control were added. The libraries were then loaded onto a flow cell and the sequencing was performed on Illumina NEXTseq.

### Quantification and statistical analysis

The quality of all of the data was checked using FastQC v0.11.5 (Andrews, 2010). The 3’ sequencing data was aligned with STAR v2.5.3a (Dobin et al., 2013) to the human genome (GRCh38.p10, Gencode comprehensive annotation). To account for the spike-in present in 4sU sequencing data, reads were aligned with STAR v2.5.3a (Dobin et al., 2013) to a combination human and yeast *Saccharomyces cerevisia*e genome (GRCh38.p10 for human and R64-1-1 for *Saccharomyces cerevisiae*). The following parameters were used: -- outSAMtype BAM --SortedByCoordinate. The counts per gene were calculated using HTseq-count (Anders et al., 2015) with the following parameters --order=pos -- stranded=no --type = exon for 3’ sequencing and --order=pos --stranded=no --type = gene for 4sU respectively. Counts and bam files for 4sU data were normalised to the spike-in prior to bigwig constructions and count plotting. Bam files were converted to count per million normalised bedgraph format using bedtools v2.30 (Quinlan and Hall, 2010), which was then converted to bigwigs using bedGraphToBigWig (Kent et al., 2010). Metagene plots were generated using deeptools (Ramirez et al., 2016) for genes separated by at least 1kb. Raw counts were analysed in R (version 4.0.5). Differential expression analysis was performed using DESeq2 package. Gene ontology terms were generated using Panther (Mi et al., 2010).

### Protein immunoprecipitation

~10 M cells were pelleted down and resuspended in 1 ml extraction buffer (20mM HEPES-Na, pH 7.4, 0,5% Triton, 600mM NaCl; in Figures 3D and S3B, extraction buffers with different concentration of NaCl or addition of urea were used, as indicated) supplemented with protease and phosphatase inhibitors (Thermo Fisher, A32961). Cell lysate was briefly sonicated 2 x 5 s (10mA, model) and centrifuged for 10 min at 14000 rpm at 4°C. The supernatant, which is a clarified protein extract, was incubated on a rotating wheel with 10 μl α-FLAG magnetic beads for 1h at 4°C. The beads were washed three times with the cold extraction buffer and during the last wash transferred to a new 1.5 ml tube. Immunoprecipitated proteins were eluted with 28 μl 1.1x LDS Sample Buffer (Thermo Fisher, NP0007) for 15 min at room temperature with vigorous shaking. After elution, 2 μl of 1 M DTT (Merck, D9779) was added and the sample was heated for 10 min at 70°C. Alternatively, proteins were eluted with 20 μl of 1 mg/ml 3xFLAG peptide (Merck, F4799) for 15 min at room temperature with vigorous shaking. After elution, 8 μl of 4x LDS and 2 μl of 1 M DTT were added and the sample was heated for 10 min at 70°C.

### SDS-PAGE and WB analyses

Proteins were separated using NuPAGE electrophoresis system (Thermo Fisher). When stained with Coomassie or silver stain, samples were run on 4-12% Bis-Tris gels (Thermo Fisher, 10247002). When analysed by western blot, samples were run on 3-8% Tris-Acetate gels (Thermo Fisher, 12095655). Prestained protein ladder (Thermo Fisher, 26619) was used as a marker. Proteins were stained with Coomassie stain prepared according to (Candiano et al., 2004) or with Pierce Silver Stain Kit (Thermo Fisher, 24612) according to the manufacturer’s protocol. For western blot, proteins were transferred from the gel to a PVDF membrane (Merck, IPFL00010) by semi-dry transfer with Trans-Blot Turbo Transfer System (Bio-Rad, 1704150) for 10 min at 25V. After transfer, membranes were blocked for 5% milk dissolved in PBST buffer supplemented with 0.05% Tween 20. Next, membranes were incubated overnight at 4°C or for 1h at room temperature with dedicated primary antibodies followed by incubation with the relevant IRDye secondary antibodies (Table S4). Detection was performed using Li-Cor Odyssey Classic Imager.

### Mass spec

The sample preparation and the mass spectrometry analysis were performed by the Advanced Mass Spectrometry Facility at the University of Birmingham.

### Cell cycle analysis

Cells were synchronised at G1/S phase using double thymidine block method. Six hours after siRNA transfection or Tet-induction, 2 mM thymidine was added to the media for 18 h, then it was removed for 9 h and added again for 15 h. Cells were washed with PBS and treated with 10 mM BrdU for 1 hour before harvesting at different time points (time 0, 4h, 8h). Cells were fixed in 70% ethanol overnight followed by 20 min incubation with 2 M HCl (containing 0.1 mg/mL pepsin) and permeabilized with PBS containing 0.2% Tween-20 and 1% goat serum. Then, cells were incubated with anti-BrdU antibody (Sigma-Aldrich) followed by incubation with FITC-conjugated secondary antibody (Sigma-Aldrich). Finally, cells were resuspended in 500 μl PI working solution (RNaseA 0.1 mg/ml, Propidium iodide 25 μg/ml) and incubated for 5-15 min at 37°C. The cell cycle analysis was done with FlowJo.

